# Imbalance of Ciliary Programs Drives Fibroblast Differentiation and Fibrotic Signaling in Systemic Sclerosis

**DOI:** 10.64898/2026.02.25.707978

**Authors:** Le My Tu Nguyen, Carlos Córdova-Fletes, Paulene Sapao, Catherine Vasquez-Hernandez, Adam Klumpp, Le Phuc Diem Nguyen, Poulami Dey, Johann E. Gudjonsson, Rebecca L Ross, Francesco Del Galdo, Natalia Riobo-Del Galdo, Priyanka Verma, John Varga, Maria E Teves

**Affiliations:** Department of Obstetrics & Gynecology, Virginia Commonwealth University, Richmond, Virginia, USA; Departamento de Bioquímica y Medicina Molecular, Facultad de Medicina, Universidad Autónoma de Nuevo León, Monterrey, México; Department of Dermatology, University of Michigan, Ann Arbor, Michigan, USA; Leeds Institute of Rheumatic and Musculoskeletal Medicine, Faculty of Medicine and Health, University of Leeds, Leeds, LS9 7TF, UK; NIHR Leeds Biomedical Research Centre, Leeds Teaching Hospitals, NHS Trust, Chapel Allerton Hospital, Leeds, UK; School of Molecular and Cellular Biology, Faculty of Biological Sciences, University of Leeds, Leeds, UK; Astbury Centre for Structural Molecular Biology, University of Leeds, Leeds, UK; Division of Rheumatology, Department of Internal Medicine, University of Michigan, Ann Arbor, Michigan, USA

**Keywords:** Systemic sclerosis, primary cilia, ciliogenesis, myofibroblast, morphogen signaling

## Abstract

Systemic sclerosis (SSc) is a chronic fibrotic disease characterized by accumulation of activated profibrotic myofibroblasts in multiple organs. The mechanisms triggering pathogenic fibroblast to myofibroblast reprogramming in SSc, and maintaining the activated myofibroblast state, are not well known. Recent studies show that primary cilia (PC), solitary sensory organelles that integrate diverse chemical and mechanical signaling pathways, may regulate fibroblast fates. Here, using an orthogonal multimodal strategy spanning human cohorts, single-cell trajectories, and targeted perturbation models, we identify a fundamental imbalance in cilia-program dynamics as a unifying driver of fibrotic activation in SSc. Meta-analysis of skin biopsies integrating eight microarray datasets and two independent scRNA-seq cohorts, revealed a conserved 15-gene cilia signature that is altered in SSc. Single-cell trajectory mapping resolved a principal progenitor→ secretory-like → myofibroblast differentiation axis, where SSc fibroblasts displayed spatially dysregulated ciliary programs that established a prolonged, disassembly-dominant “cilia-off” state. Notably, this ciliary imbalance appeared to emerge before the transition to fibrotic gene programs, positioning cilia disruption as an initiating, not secondary, event in fibroblast programming. Consistent with these transcriptomic findings, PC length was significantly reduced in SSc skin and cultured fibroblasts. Mechanistically, genetic and pharmacological perturbation studies demonstrated that cilia disruption is sufficient to drive fibrotic programs. These data establish a reciprocal regulatory framework in which shortened cilia promote sustained activation of TGF-β–Hippo feed-forward signaling, driving fibroblast-to-myofibroblast transition and amplifying fibrotic responses. Together, these findings position the imbalance between ciliary assembly and disassembly as a central determinant of fibrotic fibroblast fate in SSc. Moreover, they indicate that therapeutically stabilizing primary cilia, “ciliotherapy”, can reset pathological fibroblast trajectories and represents a promising antifibrotic strategy for SSc and other fibrotic conditions.

**Teaser:** Disrupted Primary Cilia Drives Pathophysiology of Systemic Sclerosis.

## INTRODUCTION

Systemic sclerosis (SSc), a devastating chronic fibrotic disease characterized by protean clinical manifestations and high mortality rates, lacks effective treatments (*1*). Fibrosis in multiple organs is the defining SSc hallmark that accounts for the disease’s high mortality. How fibrosis emerges synchronously in multiple distant organs in SSc remains an unanswered question with fundamental treatment implications. Myofibroblasts have emerged as the central drivers of fibrosis across different organs and conditions (*2*). A crucial concept in fibrosis is mesenchymal cell plasticity, wherein quiescent tissue-resident cells undergo activation and transform into myofibroblasts responsible for extracellular matrix (ECM) accumulation and fibrosis. In pathological fibrosis, the myofibroblasts accumulating in target organs originate from a variety of mesenchymal progenitor cells through bidirectional differentiation programs (*3*). In this process, cell-type-specific molecules for each transition are required. Well-studied signaling pathways, including TGF-β and Hippo, have been implicated in triggering and/or sustaining pathological myofibroblast transition (*4*, *5*, *6*). However, the specific mechanisms governing myofibroblast transformation and sustained activation in SSc remain elusive, representing a substantial knowledge gap and an impediment to the development of effective treatment. Our goal is to bridge this gap by elucidating the cellular and molecular mechanisms driving fibrosis and uncovering entirely novel therapeutic strategies for SSc.

Recently exciting discoveries on the potential involvement of primary cilia (PC) in the pathogenesis of SSc were made (*7*, *8*, *9*). Primary (non-motile) cilia are specialized unitary membrane organelles present in nearly all nucleated mammalian cells, and play fundamental roles in cell signaling (*10*, *11*, *12*). Acting as sensory antennae for chemical and mechanical signals, PC exert a profound influence on tissue repair and regeneration (*13*). The primary cilium is formed during G1 in the cell cycle and disassembles at the G2/M transition. Following the completion of the cell division, the cilium reassembles in G1. This cycle is finely regulated at multiple levels (*7*).

Primary cilia serve as central processing units for multiple signaling pathways, and have emerged as key hubs orchestrating cellular signaling, (*14*). Their unique structure is enriched in receptors and signal transducers regulated by a dynamic trafficking and spatial localization of proteins within the ciliary compartment. This compartmentalization generates distinct molecular signaling signatures, enabling precise spatial and temporal control of cellular responses and providing critical insight into how signaling outputs are fine-tuned (*15*). Some of these events are regulated within specific sub-compartments of the primary cilium and are influenced by the differentiation state and microenvironment of the cells in which they reside (*7*). Despite their established importance in health and disease, the mechanisms linking PC with fibrosis are not understood. Notably, key components of profibrotic signaling pathways—including TGF-β and Hippo—localize to and are regulated through the primary cilium (*11*, *16*, *17*). However, previous studies have yielded inconsistent results, with some showing increased PC length, and elevated expression of ciliogenesis-related genes, in cardiac and pulmonary fibrosis (*18*, *19*, *20*), whereas ciliopathies characterized by abnormal PC formation and function display marked myofibroblasts accumulation and fibrosis (*7*). Moreover, disruption of the primary ciliary gene *Ift88* in the mouse results in myofibroblast transition in endothelial and hepatic stellate cells (*21*, *22*). Similarly, disruption of the cilia gene *Spag17*, has been associated with myofibroblast transition in SSc (*23*). In addition, shorter PC seems to be an early feature of various fibrotic conditions (*9*). Together, these observations implicate PC in the regulation of tissue remodeling via cell-type-specific and potentially biphasic mechanisms (*7*, *24*). Indeed, we and others recently showed that in fibroblasts and endothelial cells, PC undergo biphasic changes during TGF-β-induced myofibroblast transition (*9*, *24*). PC initially appear to be required to sense TGF-β; however, as the transition to myofibroblasts unfolds, cells progressively lose PC (*24*).

Here, we further investigated the association of PC with the pathophysiology of SSc and uncovered evidence that positions PC as a previously underappreciated yet central regulator of this disease. Our findings suggest that PC critically influences myofibroblast transition and profibrotic signaling—key drivers of tissue fibrosis in SSc. This work highlights a novel mechanistic pathway with direct clinical relevance and supports the concept that therapeutic modulation of PC (ciliotherapy) may offer a transformative strategy to prevent, halt, or potentially reverse fibrosis in patients with SSc.

## RESULTS

### Reduced length of the primary cilia is a distinctive hallmark of SSc fibroblasts within the fibrotic tissue microenvironment

The size and shape of PC exert strong influence on cellular signaling and performance (*25*, *26*). Recently the association of shorter PC with fibrotic fibroblasts has been reported in various fibrotic conditions (*9*). While these *in vitro* observations provide valuable insights, a crucial step towards understanding the clinical relevance of PC dysregulation in SSc involves direct validation within the native tissue environment. Therefore, we assessed PC length directly in skin biopsies obtained from SSc patients (n=5) and healthy controls (n=4). Histological sections from these biopsies were immunolabeled with anti-ARL13B (as a PC marker) and anti-vimentin (a fibroblast marker) antibodies to enable precise visualization and measurement within the fibroblast population. Our analysis revealed reduced PC length in SSc biopsies (Fig. 1A). Consistently, dermal fibroblasts isolated from SSc skin biopsies also show shorter PC and are accompanied with increased expression of alpha-smooth actin (ASMA) when compared to those from healthy controls (Fig. 1B). These findings provide compelling evidence that shorter PC are a characteristic feature of SSc fibroblasts, whether they are cultured *in vitro* or reside within the fibrotic tissue microenvironment.

**Figure 1:**
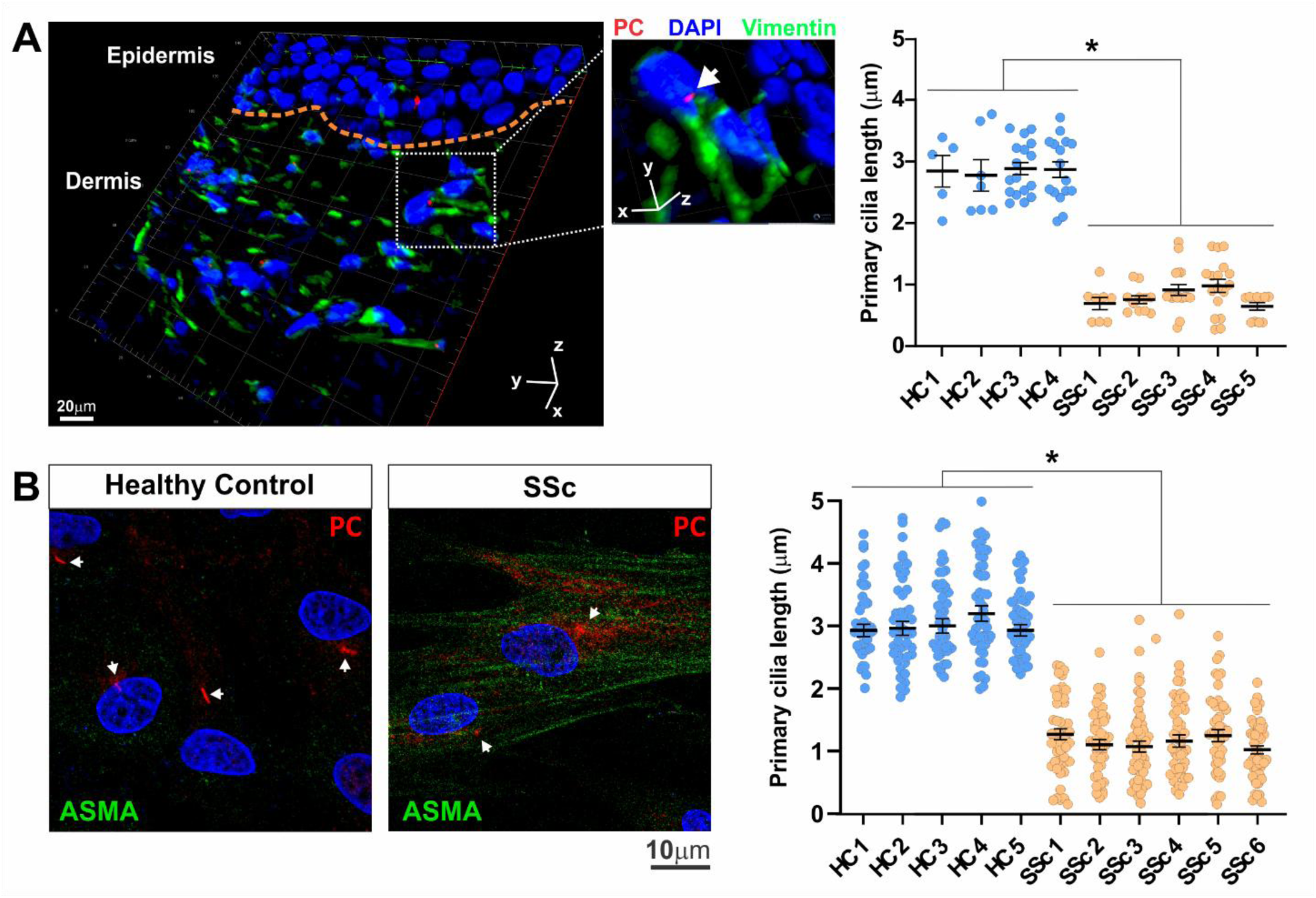
Shorter primary cilia are a distinctive hallmark of SSc fibroblasts. (A) Skin biopsies from patients with SSc (n = 5) and healthy controls (n = 4) were immunolabeled with anti-ARL13B (primary cilium marker) and anti-vimentin (fibroblast marker) antibodies to visualize primary cilia (PC) in dermal fibroblasts. Representative 3D image showing primary cilia (red) in dermal fibroblasts (green). Magnified image highlighting a fibroblast with a primary cilium (white arrow) is shown in the middle panel. Graph shows quantification of primary cilia length in SSc and healthy control biopsies. (B) Explanted fibroblasts from SSc (n = 6) and healthy controls (n = 5) were cultured and immunolabeled with antibodies against α-smooth muscle actin (ASMA) and ARL13B. A total of 50 cells per sample were analyzed. Representative images and quantification of primary cilia length are shown; arrows indicate shortened primary cilia in SSc fibroblasts. Statistical significance was assessed using One-Way ANOVA. Data are presented as mean ± SEM; * indicates statistical significance, p < 0.05.

### Shortening of the PC in SSc is governed by differential expression of cilia-related genes

#### a) Multi-platform identification of a ciliary program associated with SSc

In order to better understand the alterations in PC length noted in SSc fibroblasts, we undertook a comprehensive investigation into the regulation of ciliary gene expression. Comparing the expression profiles of SSc and healthy skin across publicly available cohorts using both bulk microarray datasets (*27*) and scRNAseq data (*28*, *29*) uncovered broad transcriptome changes indicative of ciliary dysfunction and resolution of cell-type-specific alterations within the heterogeneous skin microenvironment.

Across eight independent microarray datasets (GSE1–GSE8), we observed a distinct cilia-related signature recurrent in SSc (Table S1). Gene Ontology (GO) analysis showed significant enrichment of terms related to cilium organization/assembly, microtubules and basal body, and tubulin/microtubule binding, with additional enrichment for basal-body docking, membrane tethering, and cytoskeleton-dependent transport (Fig. 2A; Fig. S1-3).

**Figure 2:**
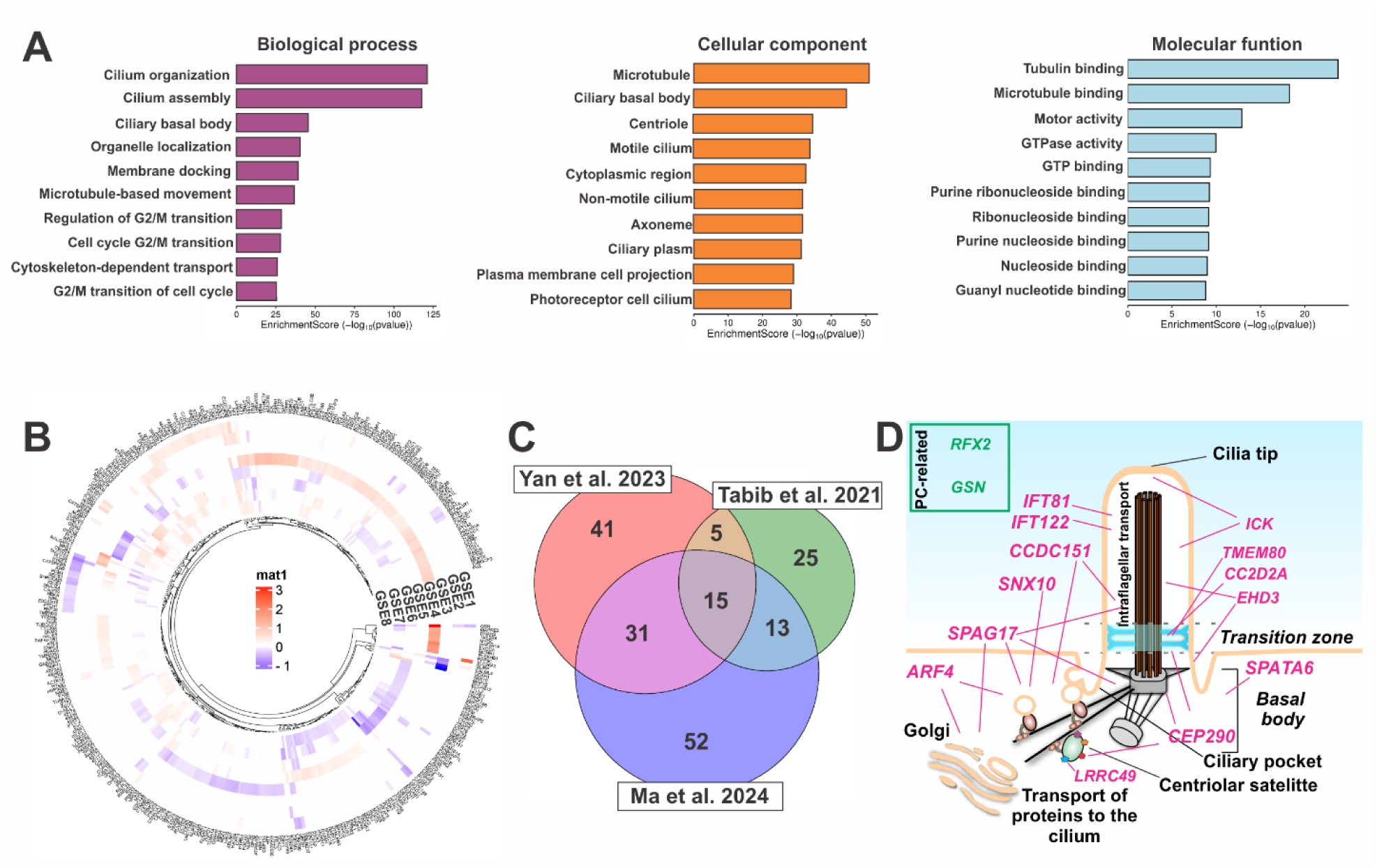
Unveiling the primary cilia signature in SSc. (A) Transcriptomic analysis of human skin microarray data from Yan et al. (2023) (*27*). Gene Ontology (GO) analysis reveals enrichment of differentially expressed genes (DEGs) associated with cilia organization/assembly and microtubule/tubulin-binding processes. (B) Circular heatmap showing cilia-related DEGs across individual GEO (GSE) datasets. (C) Venn diagram illustrating the number of significantly regulated cilia assembly/disassembly DEGs identified in Yan et al. (2023) (*27*), Tabib et al. (2024) (*28*), and Ma et al. (2024) (*29*). (D) Schematic representation of 15 cilia signature genes and their known subcellular ciliary localization.

A differential expression pattern of cilia genes as well as cilia-associated genes were represented throughout all the microarray datasets. The circular heatmap (Fig. 2B) illustrates a global overview of gene regulation, where each segment of the circle represents an individual gene, and color intensity indicates the magnitude and direction of its expression changes. The accompanying dendrogram and unsupervised hierarchical clustering heatmap further delineate relationships between gene expression profiles, representing a direct and unbiased comparison of gene regulation across the various datasets.

#### b) Single-cell resolution highlights fibroblast-centric dysregulation and a hierarchized trajectory

Next, we undertook an analysis of two independent scRNA-seq cohorts (*28*, *29*). Cross-study comparison identified differentially expressed genes (DEGs) in each dataset with a reproducible shared subset (Table S2). Analysis of GO enrichments in these single cell-level data mirrored those observed in bulk analyses (Fig. S4). Because aberrant regulation of cilia assembly/disassembly dynamic process can lead to altered ciliary length and signaling and contribute significantly to disease pathogenesis (*30*), we focused on fibroblasts and on genes that are implicated in either ciliary assembly or disassembly. We defined a 15-gene signature shared across the three data sources, encompassing core ciliary genes as well as cilia-associated genes involved in ciliogenesis, vesicle–protein trafficking, intraflagellar transport, disassembly sentinels, and transcriptional regulation (Fig. 2C-D; Table 1).

**Table 1:**
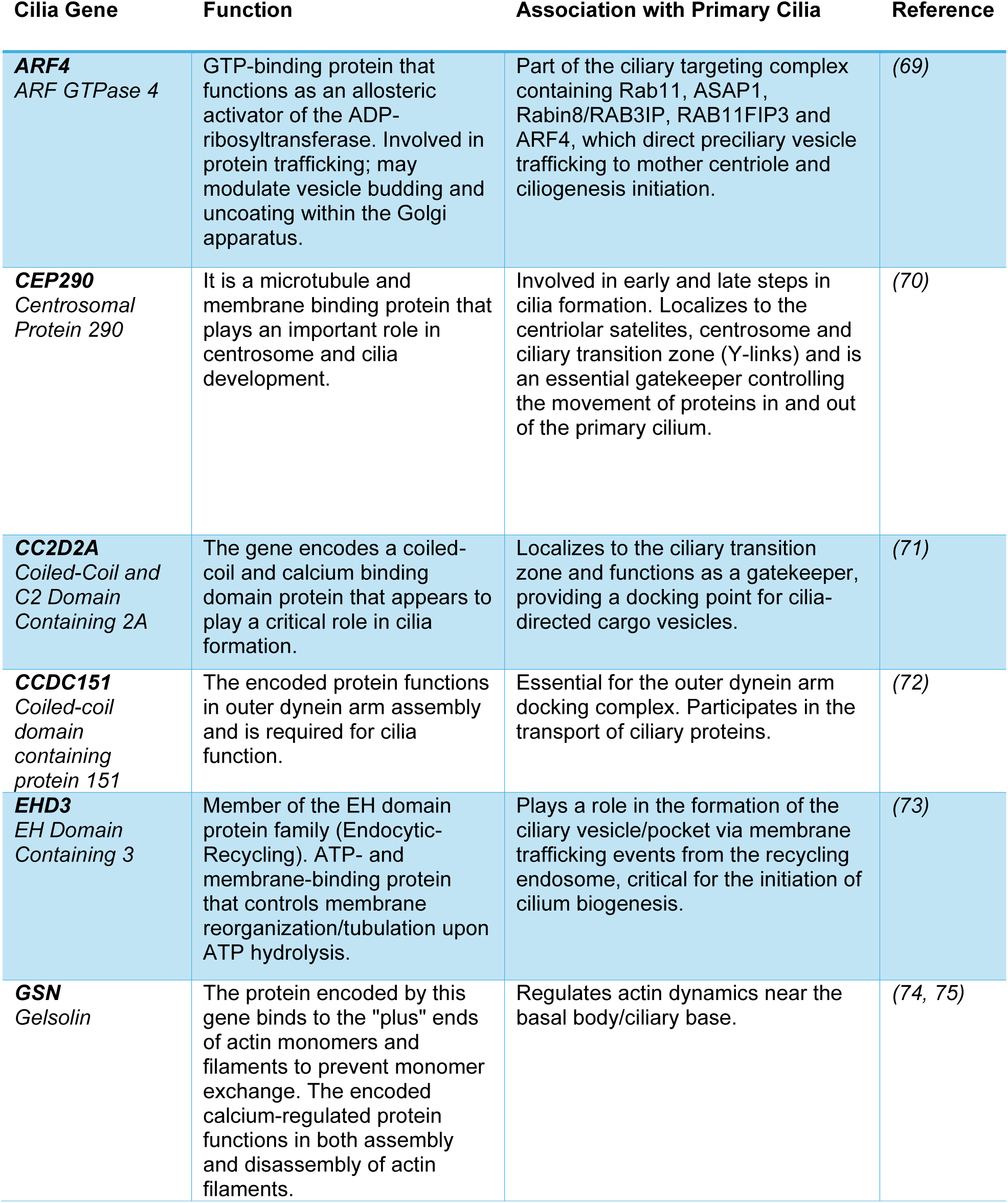
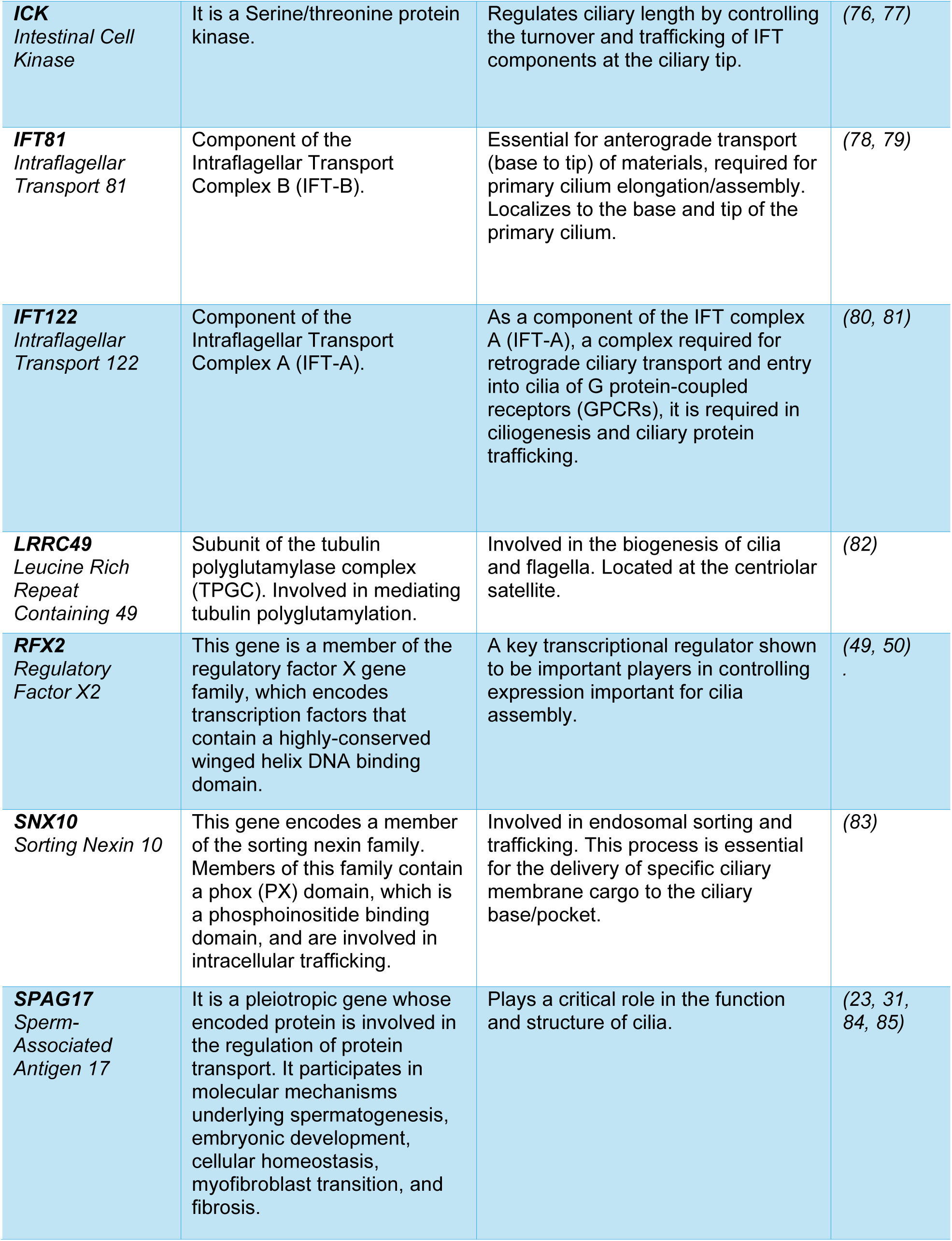
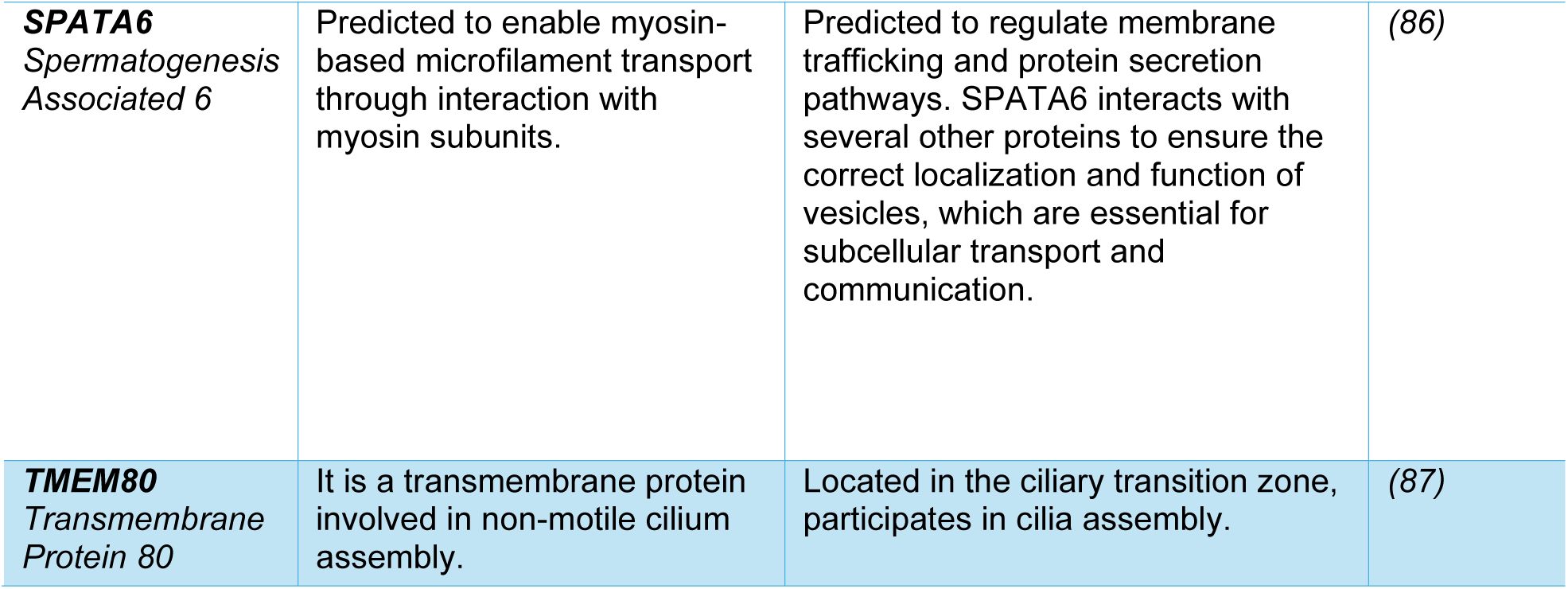
Primary Cilia Signature in Systemic Sclerosis. Fifteen cilia and cilia-associated genes were identified from transcriptomic analyses of three independent cohorts (*27*, *28*, *29*).

#### c) A unified Progenitor–Secretory-like–Myofibroblast trajectory reveals a ciliary dynamics imbalance in SSc fibroblasts

To capture dynamic transcriptional changes underlying fibroblast differentiation, we reconstructed a pseudotime trajectory spanning progenitor to myofibroblast states using our public data reported in Ma and colleagues (*29*). By dividing this trajectory into ten deciles (D1–D10), we established discrete points along the differentiation continuum that facilitated quantitative assessment of ciliary and fibrotic programs across equivalent trajectory positions. Cell counts per decile for healthy controls (n = 18) and SSc (n = 22) biopsies along the inferred differentiation path were determined, revealing that most healthy cells are distributed in early deciles, whereas SSc cells are enriched in mid and late deciles (Fig. 3A).

**Figure 3:**
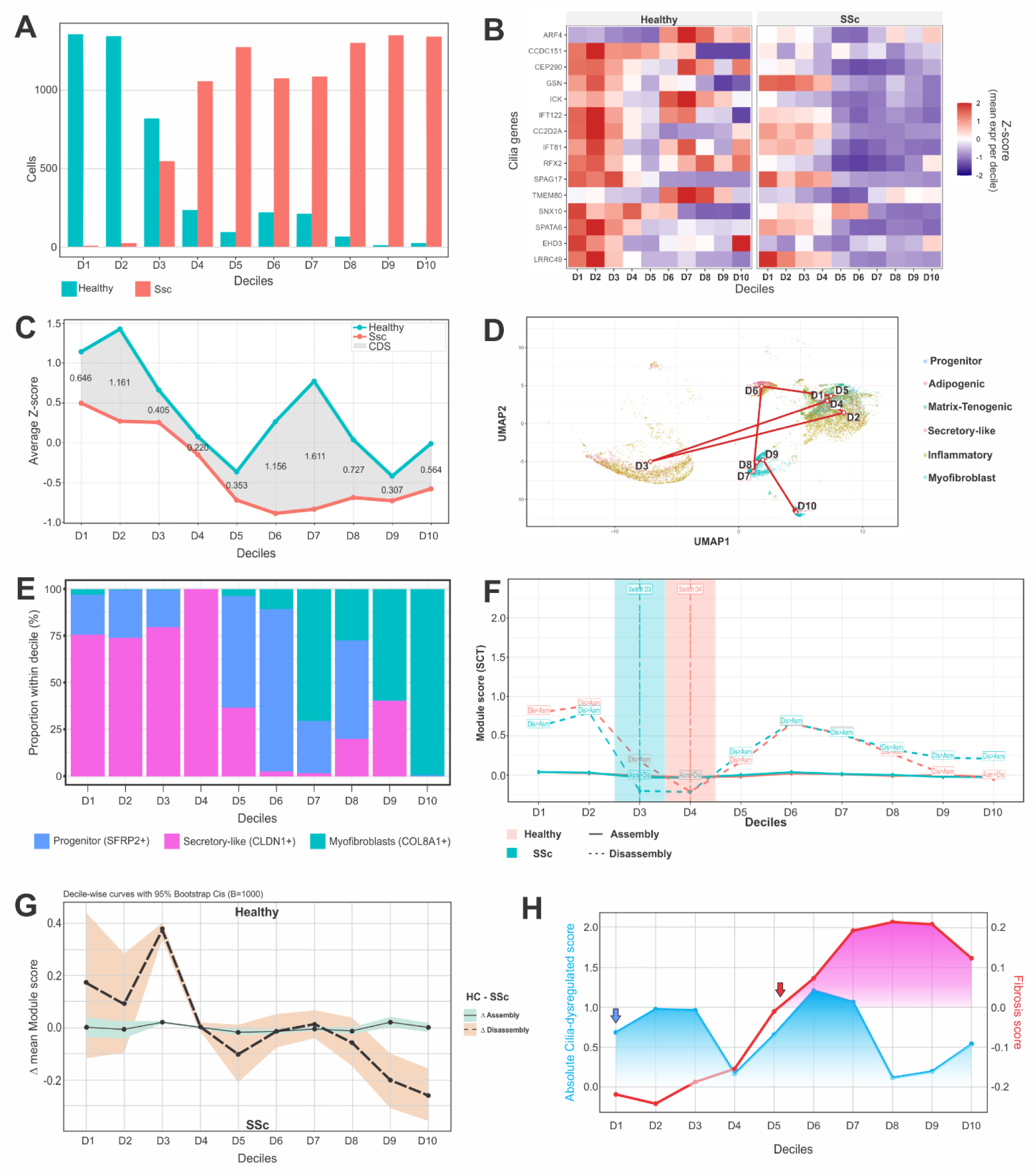
Primary cilia signature drives systemic sclerosis pathogenesis. (A) Cell counts per decile for healthy and SSc samples among trajectory cells (defined by pt_concat_P_S_M_FILTERED); deciles (D1–D10) represent type-8 quantiles of the concatenated P→S→M pseudotime. (B) Heatmap depicting cilia gene activity across fibroblast subtype clusters. (C) Visual trajectory for cilia dysregulation. The blue line represents the Healthy trajectory, the red line represents the SSc trajectory, and the shaded gray area represents the Cilia-Dysregulation Score over pseudotime deciles. (D) UMAP visualization of the inferred trajectory with decile medoids and AssemblyIndex overlay. The main progenitor→ secretory-like→ myofibroblast (P→S→M) trajectory is shown by connecting decile medoids (red path; beads D1–D10). (E) Fibroblast subtype composition restricted to the main lineage (progenitor, secretory-like, and myofibroblast) and re-normalized within each decile. (F) Cilia program dynamics along the concatenated fibroblast trajectory (healthy vs. SSc). Mean module scores for cilia assembly (solid lines) and disassembly (dashed lines) are plotted across deciles (D1–D10) along the unified P→S→M path. Labels (“Asm > Dis” and “Dis > Asm”) indicate the dominant program for each decile and condition. Vertical dashed lines denote the bootstrap-defined switch decile (first decile with mean assembly score exceeding disassembly score), with 95% confidence intervals. (G) Decile-wise differences in mean program scores are shown as ΔAssembly = mean (Assembly_Healthy) – mean (Assembly_SSc) (solid line with green ribbon) and ΔDisassembly = mean (Disassembly_Healthy) – mean (Disassembly_SSc) (dashed line with orange ribbon). Shaded bands represent 95% bootstrap confidence intervals (B = 1,000; resampling cells within each decile × condition). Values > 0 indicate higher program activity in healthy fibroblasts, whereas values < 0 indicate higher activity in SSc fibroblasts. (H) Dual-axis plot of the Abs cilia-dysregulated score and fibrosis score across pseudotime deciles, illustrating the progression of ciliary and fibrotic programs in fibroblasts along the inferred differentiation trajectory. Arrows indicate the initiation of cilia dysregulation (blue) and fibrosis burden (red).

The activity of the 15 genes identified as the cilia signature was then evaluated per decile and condition. The heatmap displays gene expression trends (Fig. 3B), revealing significant differences in cilia signature gene expression between healthy and SSc biopsies. To better quantify these alterations, we defined a cilia-dysregulated score (CDS) as the mean difference in Z-scored expression of curated cilia genes between the entire population of healthy and SSc fibroblasts across pseudotime deciles. This analysis revealed sustained ciliary transcriptional divergence beginning at the earliest pseudotime points and persisting throughout the pseudotime path (Fig. 3C), with a total integrated CDS of 7.15.

Using Monocle3 principal graphs with endpoint constraints and a signature-based corridor filter, we traced cell paths in both healthy and SSc fibroblasts and identified Progenitor (SFRP2+), Secretory-like (CLDN1+), Inflammatory (CCL19+), Adipogenic (FMO1+/FMO2+), Matrix/Tenogenic (TNN), and Myofibroblast (COL8A1+) fibroblast subpopulations, consistent with previous reports (*29*). Next, to visualize trajectory geography, we overlaid arrowed decile medoids onto a uniform manifold approximation and projection (UMAP) plot. Figure 3D shows the main Progenitor-to-Myofibroblast path in red, progressing through labeled beads D1→D10 and terminating within the Myofibroblast territory. Bead positions anchor the early–mid deciles near the Secretory-like/Inflammatory interface and the late deciles within Myofibroblast space, providing a geographic scaffold for program dynamics.

We next focused on a unified Progenitor→Secretory-like→Myofibroblast trajectory and quantified the proportion of these cells across deciles (Fig. 3E). After this cellular selection, module scoring of the 15 cilia signature genes was performed along trajectory deciles to quantify dynamic changes in ciliary activity. This analysis revealed a biphasic dynamic interplay between cilia assembly and disassembly programs. Healthy and SSc fibroblasts followed similar trajectories during early stages (D1–D2); however, SSc fibroblasts entered an assembly phase earlier (switch = D3; 95% CI D3–D3) than healthy fibroblasts (D4; 95% CI D4–D4), deviating transiently before converging again at the assembly step in D4. Following this phase, both groups transitioned into a second disassembly state. Notably, while healthy fibroblasts subsequently returned to assembly, SSc fibroblasts remained predominantly in a disassembly state through late deciles (Fig. 3F), coinciding with enrichment of myofibroblasts (which display shorter PC).

To directly compare cilia programs between conditions, we computed decile-wise differences in module scores (Δ = Healthy − SSc) for the assembly and disassembly modules along the P→S→M trajectory (Fig. 3G). ΔAssembly remained mostly positive across all deciles, indicating a persistent and maximal deficit in cilia assembly in SSc fibroblasts during the progenitor-to-myofibroblast transition. ΔDisassembly was mostly positive in early deciles but showed a pronounced decline in mid-trajectory decline (D4), corresponding to increased disassembly in SSc fibroblasts toward the myofibroblast endpoint. Consistently, SSc fibroblasts displayed fewer assembly-dominant bins, a lower peak assembly index, and an increased area under the disassembly trajectory, resulting in prolonged disassembly dominance. Collectively, these results indicate that SSc fibroblasts spend a greater proportion of their differentiation trajectory in a “cilia-off” state.

To determine whether the distinct ciliary signature we identified is mechanistically linked to primary cilia shortening and contributes to SSc pathophysiology, we next asked whether cilia dysregulation precedes and potentially drives fibroblast activation and differentiation toward the myofibroblast state. To address this, we quantified an absolute cilia-dysregulated score (CDS) and, in parallel, assessed expression of fibrosis-associated genes (COL1A1, COL1A2, COL3A1, COL5A1, COL5A2, COL8A1, FN1, ACTA2, TAGLN, CTGF, THBS1, SERPINE1). From these, we computed a fibrosis burden score across pseudotime deciles along the Progenitor→Secretory-like→Myofibroblast trajectory (Fig. 3H). Strikingly, fibroblasts in the earliest deciles already displayed significant cilia-dysregulated scores, whereas the fibrotic burden remained minimal, consistent with low extracellular matrix production and absence of myofibroblast activation at these deciles. Beyond D4, the fibrosis burden score increased sharply, reaching z-score values over zero after D5 remaining high to late deciles. These dynamics indicate that ciliary dysfunction emerges early and persists across differentiation, preceding overt fibrotic programming.

To ensure that this temporal relationship was not an artifact of the selected Progenitor→Secretory-like→Myofibroblast corridor, we repeated the analysis using the constructed pseudotime trajectory anchored on SFRP2+ healthy and SSc fibroblasts and COL8A1+ SSc myofibroblasts, as described in Ma and colleagues (*29*). This approach reproduced the same pattern, with cilia dysregulation preceding acquisition of a fibrotic transcriptomic profile (Fig. S5A).

We next asked whether this sequence of events generalizes across cohorts. Analysis of an independent scRNAseq dataset from Tabib et al. (*28*) revealed similar pattern: early deciles were dominated by cilia dysregulation, whereas fibrotic burden increased only at later pseudotime positions (Fig. S5B).

Together, these results provide quantitative and cross-cohort evidence that ciliary dysfunction in SSc fibroblasts precedes myofibroblast differentiation and fibrotic gene activation. This temporal ordering supports a model in which primary cilia disruption is not a downstream consequence of fibrosis but rather an upstream driver of pathological fibroblast reprogramming and fibrotic disease progression.

#### d) Regulators of cilia signature

To identify contextual factors influencing cilia-program activity, we computed partial Spearman correlations between the most significantly DEG signatures in SSc (genes within the GO terms for fibrotic ECM/complement module: DCN, FBLN1, COL6A2, CFD, CFH; and GO terms for cell differentiation/ cell–cell adhesion/ regulation of cytoskeleton organization/ cell cycle/ apoptosis regulation: KRT1, DSG1, DSC3, PKP1, DMKN, SFN, S100A6, PERP and NRARP (Table S3) and the 15-gene cilia panel, controlling for path position (modeled with a degree ≤3 polynomial) and condition. The top four cilia-associated targets, selected based on maximal significant |ρ| values from the frozen analysis, revealed the strongest residual relationships. ECM/complement genes showed positive correlations with GSN, whereas the other group of genes exhibited negative correlations with GSN (Fig. 4A). Edge robustness was evaluated using a condition-stratified bootstrap (B=200), and stability scores were calculated as the fraction of resamples achieving FDR < 0.05 and |ρ| ≥ 0.25. A complete stability heatmap for all 15 cilia genes and the regulators is provided in Fig. S6.

**Figure 4:**
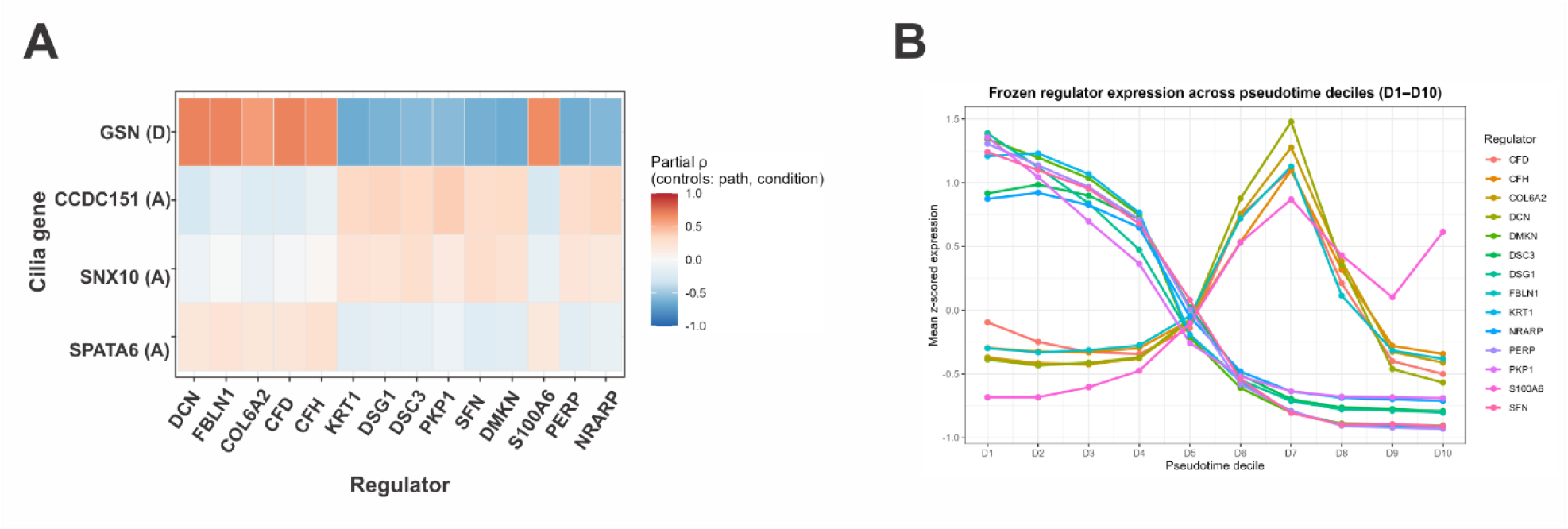
Condition-aware regulator–cilia map. (A) Heatmap showing correlations between highly scored cilia genes and cilia regulators. Partial Spearman correlations (ρ) were computed between a frozen regulator set (ECM/complement and epithelial/cytoskeletal programs) and cilia genes, controlling for path position (modeled with a degree ≤3 polynomial) and condition. Tiles represent correlation coefficients (ρ; −1 to 1). The four cilia genes displayed were selected based on the maximal significant |ρ| values from the frozen analysis. (B) Regulator expression across pseudotime deciles. Mean z-scored expression of 14 frozen regulators across pseudotime deciles (D1–D10).

Next, we evaluated the expression of the 14 regulators across pseudotime deciles to identify when do they play their highest influence (Fig. 4B). Results show that genes associated with cell differentiation/ cell–cell adhesion/ regulation of cytoskeleton organization/ cell cycle/ apoptosis regulation have an effect on cilia genes at early and middle deciles, while genes associated with fibrotic ECM/complement module have an influence in middle-late deciles with a peak at D7. To test whether the inferred regulators were robust and not due to technical artefacts, we repeated the analysis on a decontaminated, singlet-only fibroblast subset obtained by applying DecontX (contamination ≤ 0.20) and scDblFinder to the fibroblast object with pseudotime/deciles. The same 14 regulators (CFD, CFH, COL6A2, DCN, DMKN, DSC3, DSG1, FBLN1, KRT1, NRARP, PERP, PKP1, S100A6, SFN) remained the top regulators that correlate with changes in cilia gene expression and fibrosis burden scores. Decile-wise correlation heatmaps and the QC-restricted panel confirmed that regulator trajectories are essentially unchanged after decontamination, arguing that these modules reflect genuine biology rather than doublets in a scRNAseq cluster or ambient RNA contamination (Fig. S7 and S8). Collectively, these findings delineate a stable regulatory landscape in which early differentiation-related cues and later ECM/complement signals sequentially shape ciliary dysfunction, supporting a model in which cilia dysregulation is context-dependent and integrated into the emerging fibrotic program.

### Disruption of PC is sufficient to drive fibrosis

To identify mechanistic links between altered PC state and myofibroblast transition, two distinct and complementary experimental approaches were employed. First, we investigated the effect of morphological PC disruption by ablation of a PC gene. Previous studies have demonstrated that the *Spag17* gene is essential for proper ciliogenesis, as its loss in fibroblasts leads to abnormally short and dysfunctional PC (*31*). Notably, *SPAG17* was found to be one of the cilia genes with differential expression in SSc skin biopsies (*23*, *32*) (Table 1). Furthermore, cross-species transcriptome analysis between *Spag17* knockout (KO) mouse-derived fibroblasts and human SSc skin fibroblasts, revealed significant similarities in differential gene expression (*23*). Therefore, we utilized dermal fibroblasts from *Spag17* KO mice as a robust genetic system to model altered PC in SSc fibroblasts. Skin fibroblasts explanted from neonate WT and KO mice were grown to confluence, incubated for 24 hour in serum-free media to induce PC development, and then immunolabeled with antibodies for ARL13B, to detect PC, and ASMA, as a marker of myofibroblasts. Quantitative analysis of PC length and ASMA fluorescence intensity was performed on 50 cells per group and averaged for each experiment (n=3, Fig. 5A). Results revealed a significant increase in ASMA+ KO fibroblasts with genetically disrupted PC integrity.

**Figure 5:**
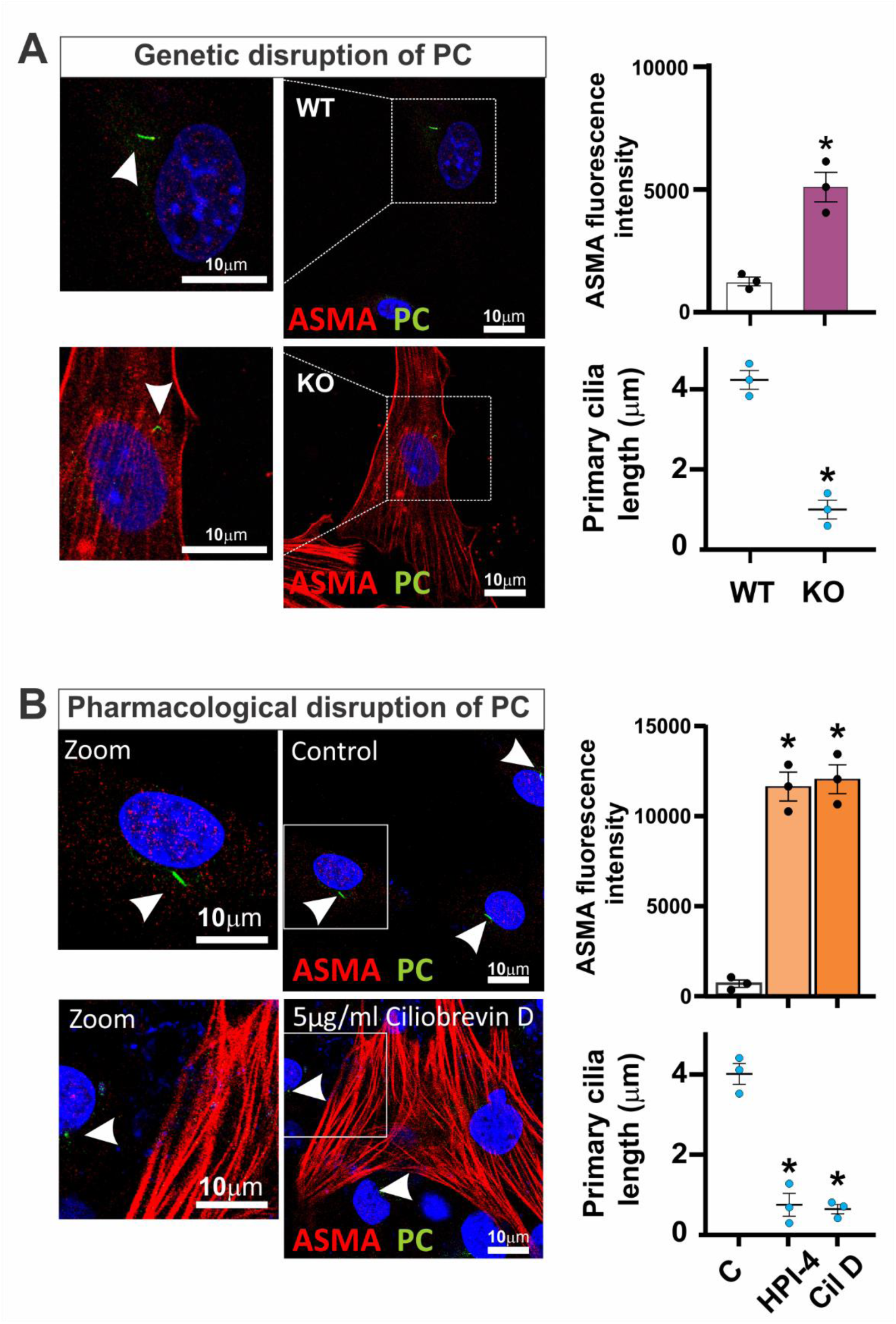
Disruption of primary cilia is sufficient to drive myofibroblast transition. Immunofluorescence analyses were performed in cultured fibroblasts using antibodies against α-smooth muscle actin (ASMA; myofibroblast marker) and ARL13B (primary cilia marker). (A) Genetic disruption of primary cilia in *Spag17* knockout (KO) fibroblasts. Representative images and quantification of ASMA expression and primary cilia length are shown for wild-type (WT) and KO fibroblasts (n=3). (B) Wild-type mouse fibroblasts were cultured in the presence or absence of the cilia inhibitors Ciliobrevin D and HPI-4 for 24 hours (n=3). Representative images are shown for control and Ciliobrevin D-treated fibroblasts; arrowheads indicate the presence of primary cilia. Graphs depict quantification of ASMA fluorescence intensity and primary cilia length. Statistical significance was assessed using Student’s t-test for two-group comparisons and one-way ANOVA for multiple comparisons. Data are presented as mean ± SEM; * indicates statistical significance, p < 0.05.

In a complementary approach to determine if pharmacological disruption of ciliogenesis would similarly induce myofibroblast differentiation, explanted WT mouse dermal fibroblasts were cultured for 24 hours in the presence or absence of well-characterized inhibitors of ciliogenesis. HPI-4 and Ciliobrevin D are small-molecule inhibitors target the AAA+ ATPase domain of the motor protein cytoplasmic dynein, impairing primary cilium assembly and leading to truncated or absent primary cilia (*33*). Following inhibitor treatment for 24 h, fibroblasts were immunolabelled for ARL13B and ASMA. The results showed that pharmacological disruption of PC was associated with significant increase in fibroblast ASMA levels, consistent with results using the genetic approach, (n=3, Fig. 5B).

Collectively, the results from both genetic models and targeted pharmacological interventions, demonstrate that disruption of primary cilia is a potent and sufficient trigger to inducer myofibroblast transition. This provides crucial functional validation for the differential expression of cilia-associated genes in SSc and strongly supports the hypothesis that dysregulation of PC dynamics plays a direct and mechanistic role in driving the fibrotic phenotype characteristic of SSc.

### Disruption of PC drives fibrosis through profibrotic TGF-β and Hippo signaling pathways regulation

Well-studied signaling pathways, including TGF-β and Hippo, have been implicated in triggering and/or sustaining pathological myofibroblast transition (*5*, *34*, *35*) and are involved in SSc pathogenesis (*29*). However, what promotes or regulates the aberrant activation of this signaling pathways in SSc fibroblast is still a missing puzzle. Primary cilia function as spatially restricted signaling hubs that integrate and regulate inputs and outputs for multiple cellular pathways (*15*). Noticeably, key regulators of both the TGF-β and Hippo pathways, including TGF-β receptors, SMAD proteins, and the Hippo effector MST1, localize to and are regulated through the ciliary compartment (*7*, *36*). On this basis, we hypothesized that the mechanism linking ciliary disruption to myofibroblast transition and fibrosis involves aberrant activation of these molecules resulting from loss of ciliary spatial control (shorter PC length).

To test this, we used nuclear accumulation of SMAD2/3 and YAP1 as readouts of TGF-β and Hippo pathway activation, respectively. Mouse dermal fibroblasts treated for 24 hours with the ciliogenesis inhibitors HPI-4 or Ciliobrevin D, as well as *Spag17*-deficient fibroblasts, exhibited a significant increase in nuclear SMAD2/3 and nuclear YAP1 (Fig. 6A and B). These findings indicate that disruption of PC length is sufficient to aberrantly activate both pathways, consistent with a model in which the cilium constrains profibrotic signaling under homeostatic conditions (*15*, *17*, *37*, *38*).

**Figure 6:**
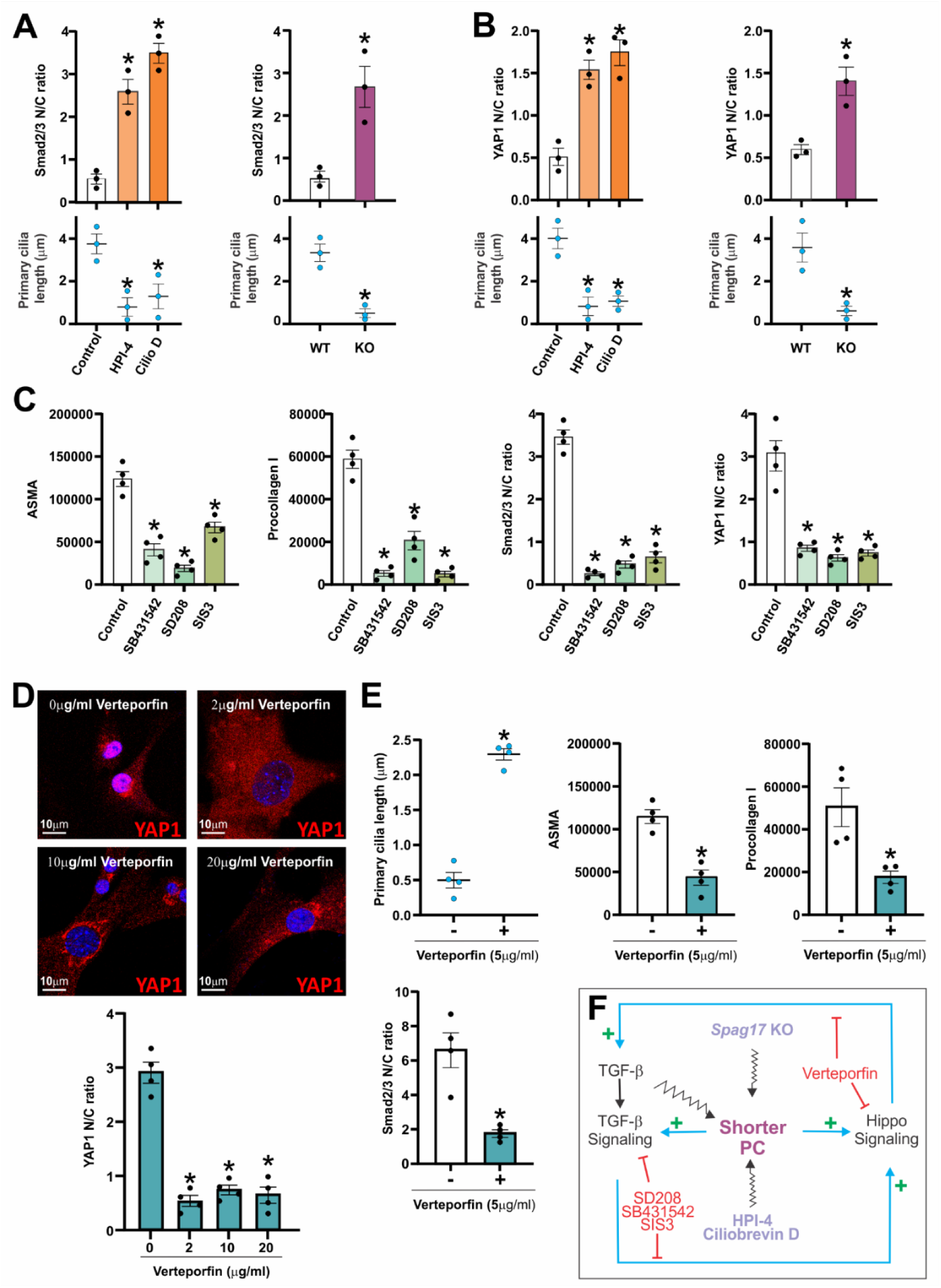
Disruption of primary cilia drives fibrosis by activating a profibrotic positive-feedback signaling loop. Pharmacological or genetic disruption of primary cilia (PC) induces activation of TGF-β and Hippo signaling pathways. Explanted wild-type (WT) mouse dermal fibroblasts were cultured in the presence or absence of the cilia inhibitors HPI-4 or Ciliobrevin D for 24 hours. Explanted *Spag17* knockout (KO) dermal fibroblasts were cultured in the absence of cilia inhibitors. Immunofluorescence analyses were then performed to assess nuclear translocation of Smad2/3 and YAP1. Fluorescence intensity in the cytoplasm and nucleus was quantified and nuclear-to-cytoplasmic (N/C) ratios and primary cilia length were calculated for each cell. Graphs show quantification of the N/C ratios of (A) Smad2/3 and (B) YAP1 from n = 3 independent experiments. (C) KO fibroblasts were cultured for 24 hours in the presence or absence of TGF-β signaling inhibitors. Graphs show quantification of immunofluorescence detection of α-smooth muscle actin (ASMA), procollagen I, Smad2/3, and YAP1 (n = 4). (D) Representative images and quantification of YAP1 immunolabeling in cultured KO dermal fibroblasts with or without Verteporfin treatment (n = 4). (E) Quantification of primary cilia length and immunolabeling for ASMA, procollagen I, and Smad2/3 in KO dermal fibroblasts after 24 hours of treatment with or without Verteporfin (n = 4). Statistical significance was assessed using Student’s t-test for two-group comparisons and one-way ANOVA for multiple comparisons. Data are presented as mean ± SEM. * indicates statistical significance, p < 0.05. (F) Schematic illustrating the mechanistic framework by which disruption of primary cilia drives fibrosis through sustained activation of a profibrotic positive-feedback loop between the TGF-β and Hippo signaling pathways. Genetic or pharmacological shortening of primary cilia promotes activation of both TGF-β and Hippo signaling. Activated TGF-β signaling further enhances Hippo pathway activation, and reciprocally, Hippo signaling reinforces TGF-β activity, establishing a self-sustaining feed-forward loop. In addition, TGF-β promotes further shortening of primary cilia, reinforcing ciliary dysfunction and perpetuating profibrotic signaling.

We next evaluated whether aberrant activation of canonical TGF-β signaling induced by disrupted primary cilia in Spag17 knockout fibroblasts could be blocked using two independent TGF-β receptor I inhibitors (10 μM SB431542 and 1 μM SD208) or by direct SMAD3 inhibition (10 μM SIS3). Incubation with each inhibitor for 24 hours significantly reduced ASMA and procollagen I levels and decreased nuclear localization of SMAD2/3 and YAP1 (Fig. 6C). These results confirm that TGF-β signaling is aberrantly activated following ciliary disruption and suggest functional cross-regulation between the TGF-β and Hippo pathways. In agreement with previous reports (*9*), TGF-β treatment of wild-type mouse fibroblasts also promoted primary cilium shortening, increased ASMA and procollagen I expression, and elevated nuclear-to-cytoplasmic ratios of SMAD2/3 and YAP1 (Fig. S9). Finally, constitutive Hippo pathway activation was reversed by treatment with Verteporfin, an inhibitor of the YAP–TEAD transcriptional complex (*39*) (Fig. 6D). Notably, treatment with 5 μg/mL Verteporfin restored primary cilium length and attenuated TGF-β pathway activation as well as ASMA and procollagen I expression (Fig. 6E).

Together, these findings establish a reciprocal regulatory framework in which shortened primary cilia promotes sustained activation of a feed-forward loop between TGF-β and Hippo pathways, driving myofibroblast differentiation and amplifying fibrotic responses. Moreover, TGF-β further promotes primary cilium shortening, reinforcing ciliary dysfunction and perpetuating a self-sustaining profibrotic signaling loop (Fig. 6F).

## DISCUSSION

### A prolonged “cilia-off” state characterizes SSc fibroblast differentiation

We provide a multi-level view of PC dysregulation in SSc. Results presented here confirm that the shortening of PC is indeed a hallmark of SSc fibroblasts, not only in vitro as previously reported (*9*), but, crucially, in native fibrotic skin. To identify the source of this phenotype, a multi-platform transcriptomic strategy encompassing several independent cohorts of SSc biopsies was used for *in silico* transcriptomic analysis. This approach identified a conserved ciliary gene signature in SSc, with genes important for cilia assembly and cilia disassembly (Table 1). Moreover, single-cell resolution, focalized in fibroblasts transitioning to myofibroblasts, revealed a pseudotime path that contemplates a unified Progenitor → Secretory-like → Myofibroblast trajectory.

A central finding of our study is a fundamental imbalance in ciliary programs. SSc fibroblasts show an earlier but attenuated transition into assembly dominance, consistent with an abortive crossing and a premature reassembly program. This is followed by a prolonged mid-trajectory disassembly-dominant phase. The net effect is that SSc fibroblasts spend a greater proportion of their differentiation path in a “cilia-off” state. The coordinated, spatiotemporal expression of assembly factors is unbalanced, leading to a disrupted cilium that is chronically targeted for resorption.

### Primary cilia are active participants in the pathophysiology of SSc

Pseudotime and decile analyses demonstrate that the imbalance in cilia programs occurs in early-intermediate fibroblast states, preceding full acquisition of *COL8A1*+, *ACTA2*+, *TAGLN*+ myofibroblast features. Moreover, our perturbation experiments showing that pharmacologic and genetic disruption of PC is sufficient to induce TGF-β and Hippo pathway activation, leading to fibroblast-to-myofibroblast transition and subsequent fibrosis. Together, these data argue that ciliary imbalance is not a late product of established fibrosis, but an active participant in the pathophysiology of SSc.

Our results are strongly supported by previous studies that show skin fibroblasts explanted from individuals with ‘very early diagnosis of systemic sclerosis’ (VEDOSS), who have no clinically detectable skin fibrosis but are at heightened risk for developing SSc within 5 years already display shorter PC comparable to that observed in SSc fibroblasts fibrosis (*9*). Moreover, fibrosis is a secondary effect of certain ciliopathies, particularly in the kidney and liver (*40*). Furthermore, several studies have implicated dysfunctional PC with the development of fibrosis in many different tissues and organs, including the heart, kidney, and lung (*41*, *42*, *43*, *44*). However, the mechanism linking PC with fibrosis may be cell type specific (*7*). This is consistent with recent spatial proteomic studies that identified intrinsic heterogeneity of PC, where 69% of the ciliary proteome is cell-type specific, and 78% exhibited single-cilium heterogeneity (*45*). The cell-type specific nature of ciliary signaling supports the need for focused studies, such as ours, to define the PC role specifically in the SSc fibroblast phenotype.

### Possible mechanisms driving cilia imbalance in SSc

Having established this altered timing and topology rooted in dual dysregulation of assembly and disassembly promoting cilia imbalance, we next sought to define the drivers of cilia imbalance. Our condition-aware regulator mapping points to the fibrotic ECM and complement as key participants. ECM/complement genes (DCN, FBLN1, COL6A2, CFD/CFH) are positively associated with GSN, the actin-severing driver of cilium disassembly, with ICK as an additional disassembly component (Fig. 4 and Fig. S6).This mechanotransduction axis is consistent with established concepts: a) Integrin/FAK activation that senses matrix stiffness and initiates signaling cascades essential for fibroblast migration and fibrosis (*46*). b) RhoA/YAP/Hippo signaling activation, which links cytoskeleton remodeling and mechanical cues to the profibrotic transcriptional output, with the Hippo pathway being a critical promoter of myofibroblast differentiation in SSc (*29*). c) AURKA/HDAC6-mediated cilium resorption, a mechanism where Aurora A kinase activates Histone Deacetylase 6 to deacetylate α-tubulin, destabilizing the ciliary axoneme and inducing disassembly (*47*).The activation of the NEDD9/AURKA/HDAC6 axis as potentially responsible for the loss of cilia in fibrosis has been previously reported (*44*), however its involvement in SSc fibrosis and how and when this mechanism is activated needs further investigation. In any case, these mechanosensing →actin remodeling →cilium disassembly axis activates after the first peak shown in the cilia-dysregulated score, suggesting it may be more related to the prolonged disassembly dominance in SSc observed at late deciles (Fig. 3G) and not the initiator of ciliary imbalance (Fig. 3H and 4B). On the other hand, genes important for cell differentiation/ cell–cell adhesion/ regulation of cytoskeleton organization/ cell cycle/ apoptosis regulation: KRT1, DSG1, DSC3, PKP1, DMKN, SFN, S100A6, PERP, NRARP, LGALS7B and FXYD3; participate at early deciles (D1-D4), suggesting they may be responsible for the basal cilia imbalance detected at early deciles in SSc fibroblasts.

Another possibility is that the altered expression of the 15 cilia signature genes in SSc fibroblasts is driven, at least in part, by upstream epigenetic mechanisms. Prior studies have shown that *SPAG17*, a gene associated with ciliogenesis, is downregulated in SSc fibroblasts in association with reduced chromatin accessibility (*23*), and additional ciliary genes have also been reported to undergo epigenetic silencing in SSc (*48*). These data suggest that chromatin remodeling may directly restrict ciliogenesis programs. In addition, RFX2, a master transcription factor that controls the expression of cilia genes (*49*, *50*), is one of the genes downregulated in SSc (Table 1), suggesting that early loss of RFX2 may modulate the transcription of several other genes within the 15-gene ciliary signature.

### Primary cilia regulation of profibrotic signaling pathways

SSc is a clinically heterogeneous fibrotic disease with no effective treatment to date. How SSc patients develop fibrosis synchronously in multiple distant organs is an unanswered question with fundamental treatment implications. Myofibroblasts have emerged as the central drivers of fibrosis across different organs (*2*). A crucial concept in fibrosis is mesenchymal cell plasticity, wherein quiescent tissue-resident cells undergo activation and transform into myofibroblasts responsible for ECM accumulation and fibrosis. In pathological fibrosis, the myofibroblasts accumulating in target organs originate from a variety of mesenchymal progenitor cells through bidirectional differentiation programs (*3*). In this process, cell-type-specific molecules for each transition are required. Well-studied signaling pathways, including TGF-β and Hippo, have been implicated in triggering and/or sustaining pathological myofibroblast transition (*5*, *34*, *35*). However, the specific mechanisms governing myofibroblast transformation and sustained activation in SSc remain elusive, representing a substantial knowledge gap and an impediment to the development of effective treatment. In recent years, attention has been turned to the role of PC in the pathophysiology of SSc, a concept that has been largely unexplored in Rheumatology until very recently (*7*). Notably, PC are specialized solitary membrane organelles present in nearly all nucleated mammalian cells, acting as sensory antennae with fundamental roles in chemical and mechanical cell signaling.

We have investigated here whether PC participation in the pathophysiology of SSc was mediated via activation of profibrotic TGF-β and Hippo signaling pathways, since key components of these signaling pathways localize to the PC (*11*, *16*, *17*). Our results show that both signaling pathways are activated as a result of shortening PC in a feed-forward loop mechanism. This is in agreement with previous studies showing interdependence of TGF-β and Hippo signaling pathways in myofibroblast differentiation (*5*, *34*, *35*) and the participation of these signaling cascades in SSc pathophysiology (*9*, *29*). Our work found that PC were the missing important piece of the puzzle linking the regulation of these signaling cascades and setting the base that PC could be the central hub coordinator of several cell signaling pathways previously reported to participate in SSc pathophysiology, including Wnt, Hedgehog (HH), Notch, PDGF, GPCR, and mTOR signaling (*51*, *52*, *53*). For instance, PC are also enriched in molecules from these signaling pathways, and their regulation and signaling has been shown to be mediated via PC (*14*, *54*). However, further investigation on how they are activated in SSc and when needs further research.

### Clinical implications and future directions

The present results have significant potential implications. The findings highlight that altered PC length can be restored (normalized) by various compounds (*55*, *56*), and modulation of PC length such as “ciliotherapy” represents a potential approach to preventing or even reversing fibrosis in SSc (*9*, *13*). Data presented here expose tractable intervention points that target the cilia dysregulation, proposing a clear and actionable path for developing disease-modifying therapies centered on ciliary integrity for SSc. We have identified five promising therapeutic strategies, beginning with *cilium stabilization* through AURKA or HDAC6 inhibition (*47*) to block ciliary resorption and extend the narrow assembly window. Several HDAC6 inhibitors are currently under study for their ability to restore cilia length and counteract aberrant signaling (*57*), suggesting a novel repurposing approach for SSc. The second strategy is targeting PC *mechanotransduction* signaling regulation, employing FAK or YAP inhibition to decouple cells from the stiff matrix and prevent ECM/ complement-driven disassembly initiation; this is already supported by studies on the YAP inhibitor, Verteporfin, as a potential treatment for scarring and fibrosis in SSc (*29*, *58*). Third, *complement modulation* involves targeting the CFD/CFH axis upstream of the disassembly cascade. Fourth, *ciliogenesis* support seeks to enhance ARF4/RFX2/IFT activity to permit re-ciliation or to target cAMP production using PDE inhibitors, such as the PDE4B inhibitor BI 1015550, which has shown antifibrotic properties for IPF (*59*), or Fenoldopam, which promotes cilia growth via Dopamine Receptor DR1 (*56*, *60*). Finally, we propose *nano-therapy* for PC-specific targeting and localized drug delivery (*61*) (Fig. 7).

**Figure 7:**
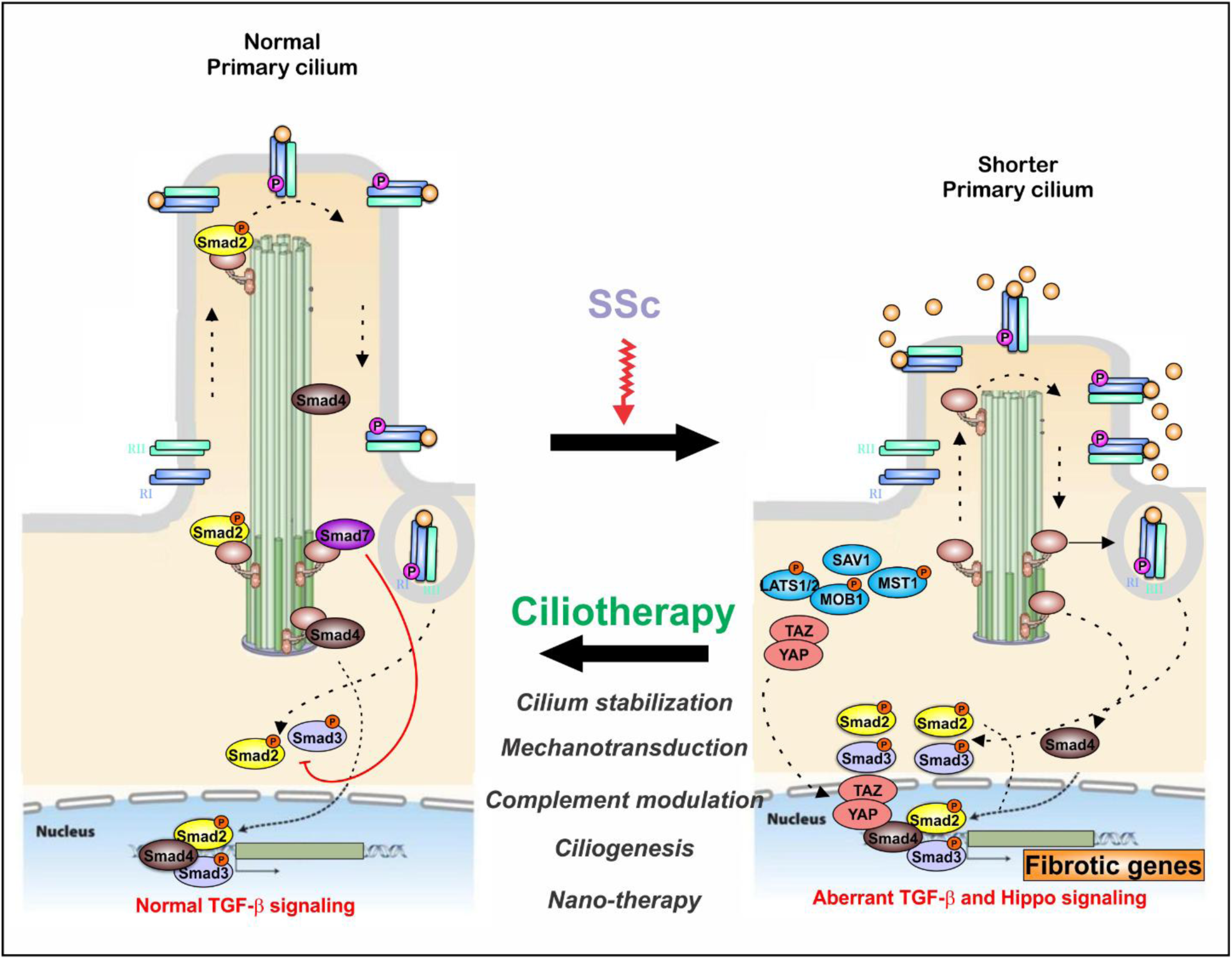
Therapeutic modulation of PC such as “ciliotherapy” represents a promising avenue for preventing or even reversing fibrosis in SSc. We have identified five promising therapeutic strategies: (i) stabilization of primary cilia through inhibition of HDAC6 or AURKA; (ii) modulation of cilia-dependent mechanotransduction signaling via inhibition of FAK or YAP; (iii) regulation of complement-driven ciliary disassembly by targeting the CFD/CFH axis upstream of the disassembly cascade; (iv) promotion of ciliogenesis using compounds that enhance re-ciliation; and (v) cilia-targeted drug delivery approaches, including nanotherapeutic strategies, to selectively modulate ciliary signaling.

In conclusion, our findings compel a paradigm shift in the understanding of SSc pathophysiology, in which disruption of the primary cilium emerges as a driver, rather than a consequence, of fibrosis. We demonstrate that an intrinsic imbalance between ciliogenesis and cilia-disassembly programs locks SSc fibroblasts into a prolonged “cilia-off” state, establishing an early event that potentiates profibrotic signaling. Thus, we propose that strategies aimed at stabilizing primary cilium structure and length (“ciliotherapy”) may have the potential to reset pathological fibroblast trajectories and ameliorate fibrosis in SSc patients.

## MATERIALS AND METHODS

### Human Subjects

Skin biopsies were taken from the affected forearm of patients. The study was approved by the University of Michigan Institutional Review Board (IRB), and all patients gave written consent. The study was conducted according to the Declaration of Helsinki Principles. Clinical information for subjects in this study was reported in Verma et al., 2025 (*9*).

### In silico RNA-seq analysis of microarrays

Key software & libraries. *In silico* analysis was performed using 10x Genomics Cell Ranger (demultiplexing/alignment/UMI quantification). R (v4.x). Seurat (v4/5) for QC, integration, SCT normalization and clustering (*62*, *63*, *64*). UMAP for embedding (*65*). Monocle3 for principal-graph learning and pseudotime (*66*, *67*). Visualization with ggplot2 (*68*), ggrepel, ggnewscale, patchwork.

Single-cell raw UMI matrices (per-sample filtered_feature_bc_matrix, Healthy and SSc) were downloaded from GEO (GSE249279), and processed using a uniform Seurat/SingleCellExperiment (SCE) pipeline. Fibroblasts were isolated, and low-quality cells, doublets, and cells with >15% mitochondrial UMIs were removed. The resulting dataset contained 63,830 high-quality fibroblasts with UMAP embeddings and Slingshot pseudotime assignments. For downstream modeling, full transcriptomic data were merged into the fibroblast object (sce_restored) to retain counts and logcounts for pseudotime lineage cells alongside all metadata.

Fibroblast subtypes were harmonized to Progenitor (SFRP2⁺), Secretory-like (CLDN1⁺), Inflammatory (CCL19⁺), Myofibroblast (COL8A1⁺), Adipogenic, and Matrix/tenogenic (stored as subtype_pretty). Assignment was validated by per-cluster marker enrichment. Trajectory structure was learned with monocle3 principal graphs (Seurat → CellDataSet conversion), and paths were traced between subtype anchors with a custom trace_path_between_subtypes wrapper enforcing endpoint constraints, graph-partition consistency (same_partition = TRUE), and a signature-based corridor filter (path_cell_filter = “signature_corridor”, corridor_q = 0.8) to suppress side-branch bleed-through. Two segments—Progenitor→Secretory-like (PS) and Secretory-like→Myofibroblast (SM)—were re-rooted to 0–1 and concatenated (PS→[0,0.5], SM→[0.5,1], averaging overlaps) to yield pt_concat_P_S_M_FILTERED; deciles D1–D10 were defined with type-8 quantiles (fallback to equal spacing if needed). For memory-light trajectory visuals, decile medoids (UMAP points nearest per-decile medians) were connected with arrows; optional beads labeled D1→D10.

Ciliary programs were scored with AddModuleScore: Assembly = {*ARF4*, *RFX2*, *IFT81*, *IFT122*, *CC2D2A*, *CCDC151*, *SPAG17*, *SPATA6*, *LRRC49*, *CEP290*, *EHD3*, *SNX10*, *TMEM80*}; Disassembly = {*ICK*, *GSN*}. The AssemblyIndex = Assembly − Disassembly was overlaid on UMAP. For figure 3F, mean program scores were summarized by decile × condition (Healthy, SSc) and plotted (Assembly solid; Disassembly dashed) with program-dominance badges; the “switch decile” (first decile where mean Assembly > Disassembly) was estimated per condition by bootstrap (B = 1,000), stratified by condition, with median and 95% CIs drawn as vertical dashed lines.

Quantitative Scoring. Cilia scores were computed as: 1) cilia-dysregulated score calculated as 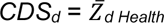 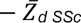, where 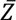 is the average Z-score of all 15 cilia genes in that decile. The Integrated Dysregulation was calculated as 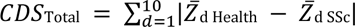 and the absolute CDS as 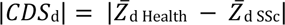. Fibrosisscore was calculated as 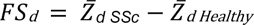, where 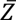 is the average Z-score of the fibrosis associated genes in that decile.

Regulator→cilia relationships were mapped with partial Spearman correlations computed on rank-transformed residuals: for cells on the concatenated path, an expression matrix (cells × genes) was built from the SCT data slot for a frozen regulator set (DCN, FBLN1, COL6A2, CFD, CFH, KRT1, DSG1, DSC3, PKP1, SFN, DMKN, S100A6, FXYD3, PERP, LGALS7B, NRARP) and the 15-gene cilia panel. Each gene was residualized against a design ∼ poly (path, degree ≤ 3) + condition (degree capped by unique path positions; condition included when >1 level), then residuals were ranked and z-scored prior to correlation. Edge robustness was assessed via bootstrap resampling (B=200). To evaluate predictive power, logistic regression and SVM (radial kernel) classifiers were trained (70/30 split) on cilia Z-scores and ECM z-score features. Two-sided p-values were derived from t-statistics and adjusted by BH-FDR; edges were reported at FDR < 0.05 and |ρ| ≥ 0.25. For reproducibility, we froze the path cell set, regulator list, and seed = 202502, exporting a top-4 cilia heatmap, a full 15-gene heatmap, and an edge-stability table from a condition-stratified bootstrap (B = 200). Figures were generated with ggplot2/patchwork and annotate the data source (human scleroderma skin; scRNA-seq ± spatial; GEO: GSE249279).

Single-cell QC robustness (keratin ↔ GSN re-check). Ambient RNA was removed with DecontX (run on raw counts when available; contamination threshold ≤ 0.20) and doublets were filtered with scDblFinder (per-sample when orig.ident was available). To test robustness of the keratin–GSN antagonism, we re-estimated partial Spearman correlations between keratin/epithelial junction regulators (KRT1, DSG1, DSC3, PKP1, DMKN, SFN) and GSN on the frozen path-cell set after (i) excluding top-1% keratin outliers, (ii) DecontX decontamination, and (iii) scDblFinder singlet-only filtering. Partial correlations were computed exactly as in the main analysis (residualized ranks after linear residuals on ∼ poly (path, degree ≤ 3) + condition, BH-FDR). All six edges remained significant and shifted modestly more negative (median Δρ ≈ −0.05), as summarized in Keratin_vs_GSN_before_after_cleaning.csv and visualized in Fig. S7 and S8.

To ensure computational reproducibility and minimize figure churn, all final analyses were performed using a fixed random seed, cell set, and regulator list and we report bootstrapped confidence intervals for switch-deciles and edge stabilities from this finalized configuration.

### Spag17 knockout fibroblasts

Dermal fibroblasts from neonate WT and KO (Spag17/Sox2-Cre) mouse line [Sapao et al., 2023] were extracted following standard procedures (*31*). Briefly, neonates were euthanized according to protocol AM10297 approved by the VCU Institutional Animal Care and Use Committee. Skin was removed from newborn mice under sterile conditions and digested by incubation with 0.25% trypsin/EDTA (Gibco, Life Technology Corporation, NY) at 37°C for 30 min and pipetting up and down every 5 minutes to mechanically assist with the tissue digestion. After digestion, cells were washed once with PBS and centrifugated for 5 min at 2,000 rpm. Supernatant was resuspended in 5 ml PBS and filtered through a 100 µm cell stainer. The filtered suspension was centrifuged for 5 min at 2,000 rpm and pellet resuspended in culture media containing DMEM (Gibco, Life Technology Corporation, NY), 10% FBS (Gibco, Life Technology Corporation, NY) without antibiotics. The cells were plated into an 8 well chamber slide (Falcon, Corning Inc., NY) and cultured until 75-90% confluence. Media change was conducted every other day. Primary cilia development was induced by serum starvation in DMEM (Gibco, Life Technology Corporation, NY), media without FBS and without antibiotics. Treatment with 10 μM SB431542; 1 μM SD208, 10 μM SIS3 or 5µg/ml Verteporfin (Sigma Aldrich, St. Louis, MO), was conducted during 24 hours in culture media without FBS, so the inhibition occurs at the same time of serum starvation. P0 or P1 passages cells were used for all the experiments.

### Immunofluorescence

At the end of the experiments, cells were fixed in 4% formalin (Sigma Aldrich, St. Louis, MO) and processed for immunolabelling (*31*) using anti Arl13B (Thermofisher, #17711-1-AP), Vimentin (Novus Biologicals, # NBP1-92687), Procollagen I (Abcam, #ABT257), ASMA (Abcam, #AB5694), Smad2/3 (Cell signaling, #8828) or YAP1 (Sigma, #HPA070359) primary antibodies overnight at 4 °C. After washing, cells were incubated with anti-rabbit Alexa Fluor 488-labeled (1:3000), anti-mouse Alexa Fluor 488-labeled (1:3000), anti-rabbit Cy3-labeled (1:5000), or anti-mouse Alexa Fluor 594-labeled (1:5000) secondary antibodies, respectively to primary antibody specie (Jackson ImmunoResearch Laboratory Inc., Grove, PA), washed with PBS and mounted with VectaMount with DAPI (Vector Laboratories, Inc., Burlingame, CA).

### Image analysis

Images were captured under a Zeiss LSM 700 confocal laser-scanning microscope and analyzed using ImageJ as previously reported (*23*, *31*). Accurate measurements of PC length were obtained by capturing 3D confocal images. Around 50 cells per treatment and experiment were analyzed for primary cilia length, fluorescence intensity and N/C ratio quantification.

### Statistical analysis

GraphPad Prism 10 software was used for statistical analysis. Data are presented as means ± standard error. Data were analyzed comparing the means from two groups by Student’s t-test or One-Way ANOVA for multiple comparisons. Means were considered significantly different when the p-value was < 0.05. Statistics for large data used analysis in R.

## ACKNOWLEDGMENT

Services in support of the research project were generated by the VCU Massey Comprehensive Cancer Center Microscopy Shared Resource, supported, in part, with funding from NIH-NCI Cancer Center Support Grant P30 CA016059.

## Funding

supported by grants from:

The National Scleroderma Foundation (MET, JV)

Leo Foundation (MET and JV)

Rheumatology Research Foundation (MET and JV)

Department of Defense (JV and MET).

NIH R01-AR069071 (JEG)

NIH R01-AR073196 (JEG)

NIH P30-AR075043 (JEG)

NIH R21 AR077741 (JEG)

Susan Cheney endowment for Scleroderma Research (FDG)

## Author contribution

Conceptualization: MET, JV

Investigation: LMTN, CC-F, PS, CV-H, AK, LPDN, PD,

Formal Analysis: MET, PD, CC-F

Resources: MET, JV, JEG

Visualization: MET, CC-F

Supervision: MET, JV

Writing—original draft: MET, CC-F

Writing—review & editing: MET, RLR, NR-DG, FDG, JV, JEG

Project administration: MET

Funding acquisition: MET, JV

## Competing interests

J.E.G. has received Grant support from Celgene/BMS, Janssen, Eli Lilly, and Almirall. J.E.G. has served on advisory boards for AstraZeneca, Sanofi, Eli Lilly, Boehringer Ingelheim, Novartis, Janssen, Almirall, BMS. All other authors declare they have no competing interests.

## Data and materials availability

All data, and materials used can be found within the article and its supplementary information.

## Supplementary Materials for

**Figure S1:**
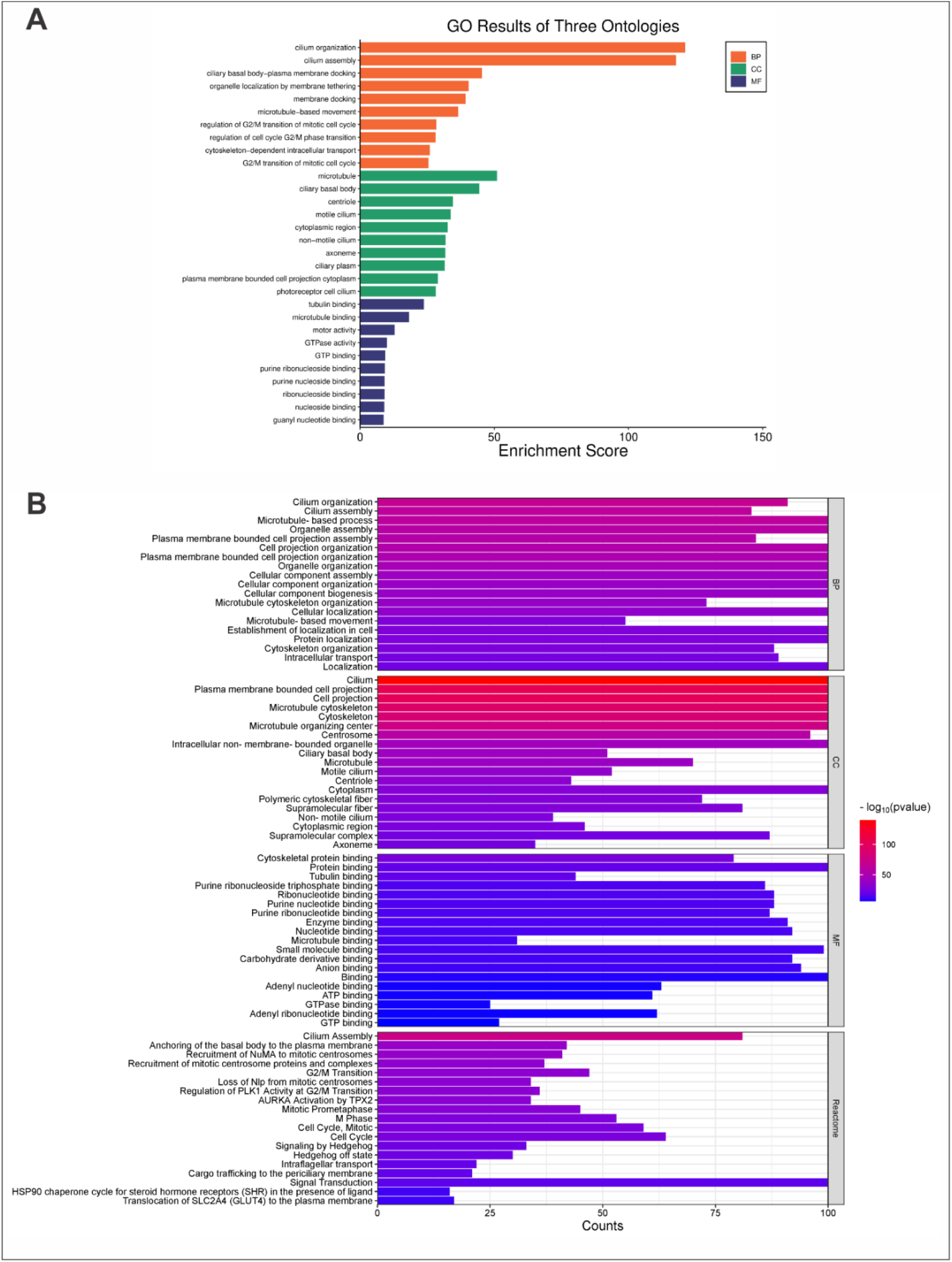
Gene Ontology analysis of biological processes, cellular components, and molecular functions in microarray datasets. (A) Bar plot showing the top Gene Ontology pathways enriched across the eight GSE microarray datasets. (B) Bar plot showing gene counts for the top Gene Ontology terms across the eight GSE microarray datasets.

**Figure S2:**
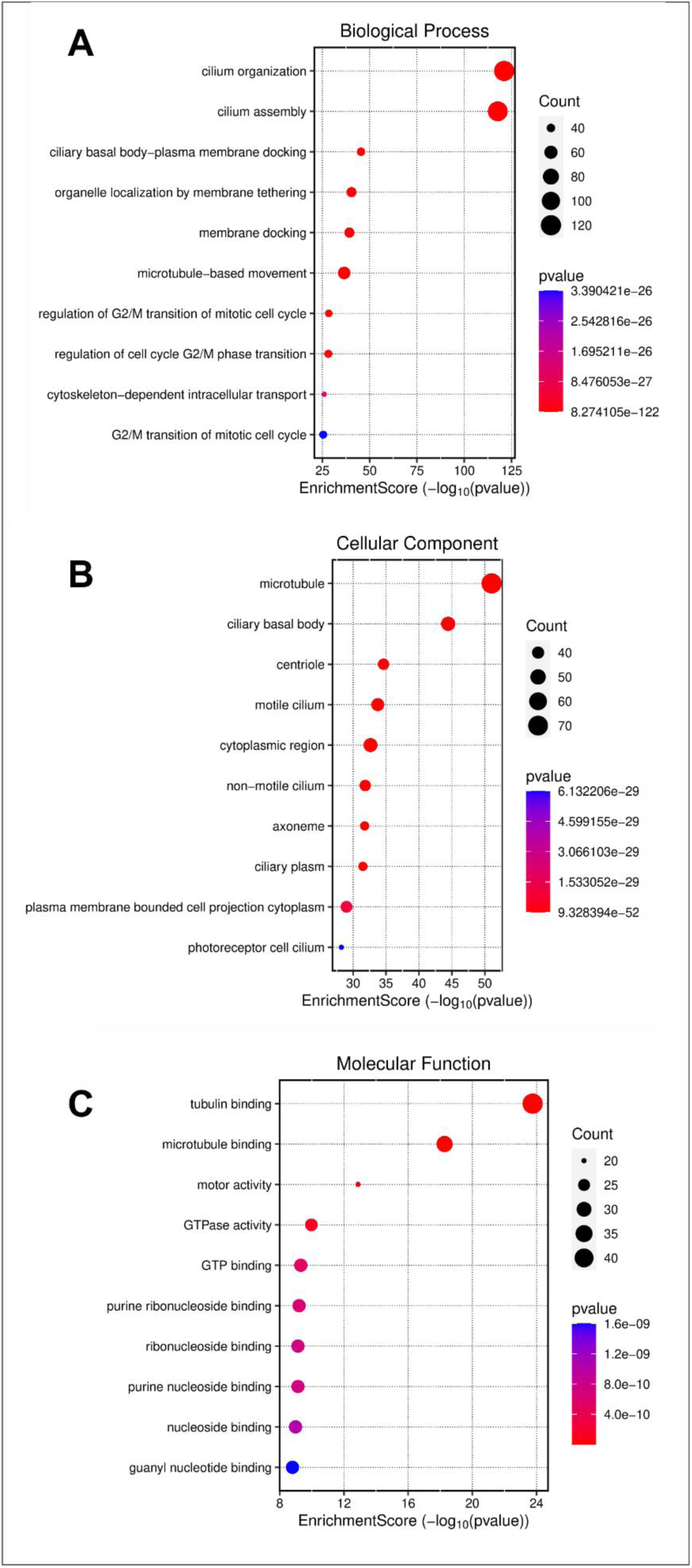
Gene Ontology analysis of biological processes, cellular components, and molecular functions in microarray datasets. Bubble plots showing enrichment scores for (A) Biological processes. (B) Cellular components. (C) Molecular functions.

**Figure S3:**
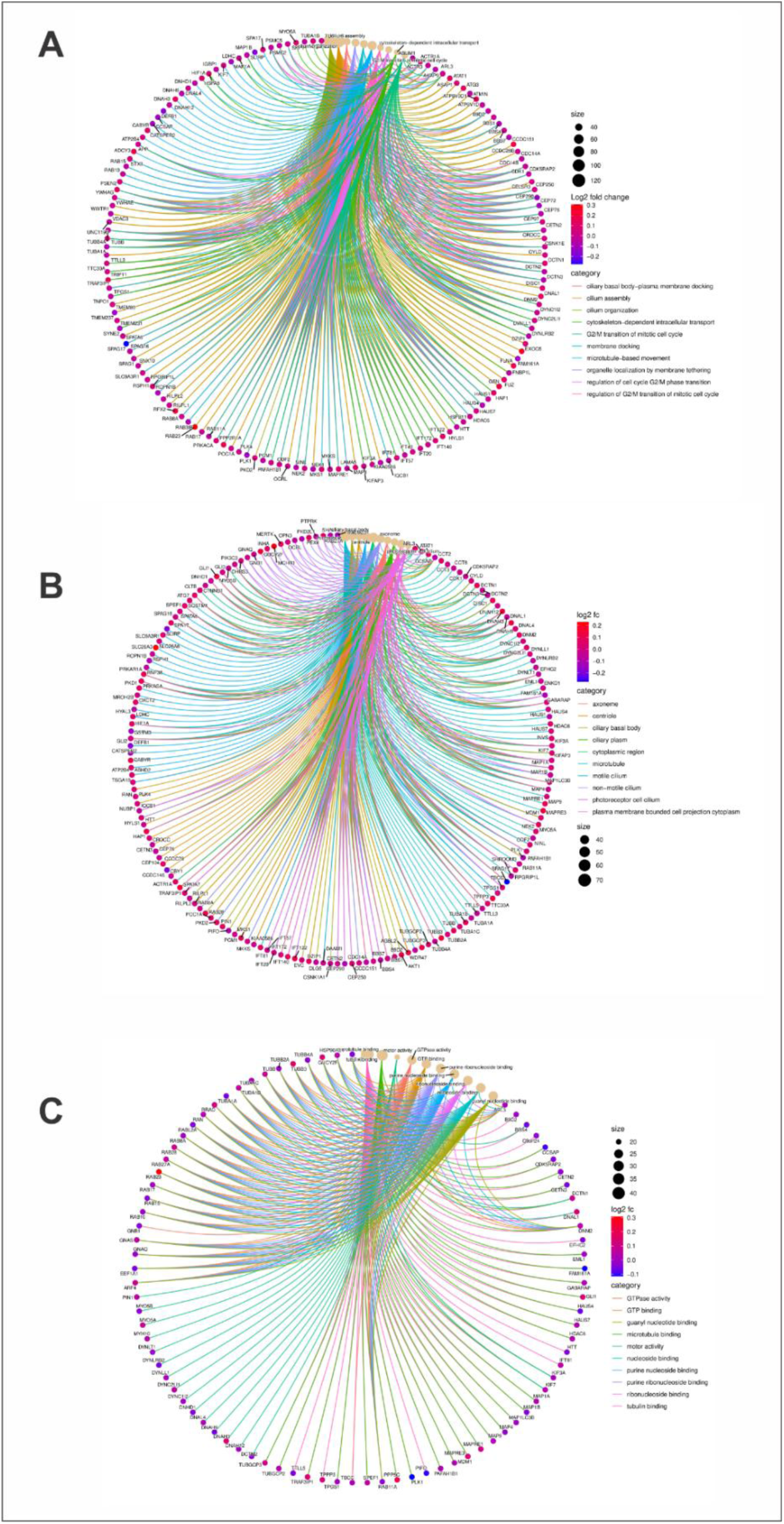
Gene Ontology analysis of biological processes, cellular components, and molecular functions in microarray datasets. Chord diagrams illustrating RNAseq analysis for (A) Biological processes. (B) Cellular components. (C) Molecular functions.

**Figure S4:**
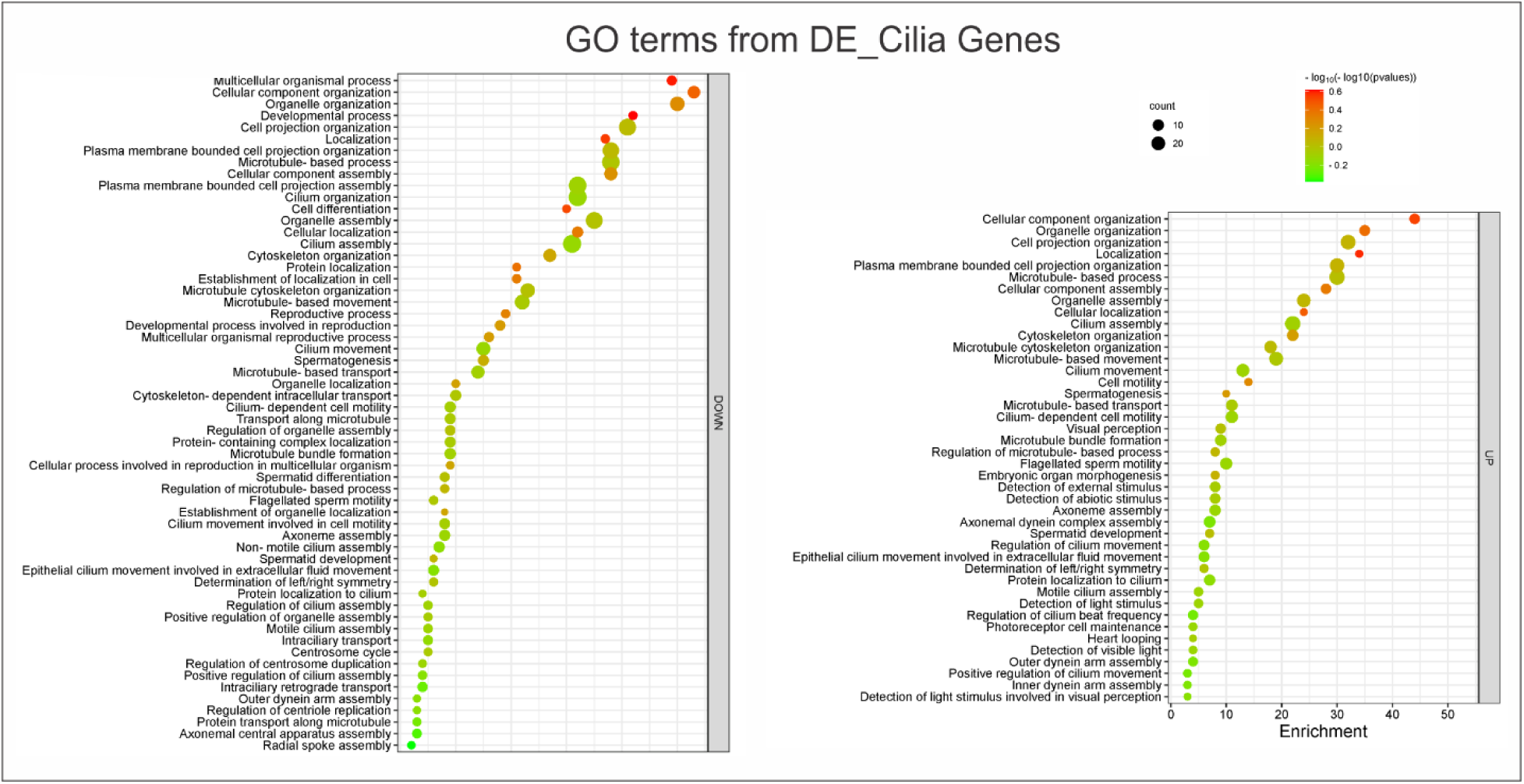
Gene Ontology analysis of the scRNA-seq dataset from Tabib et al. (2021) (28). Bubble plots showing the top enriched Gene Ontology pathways.

**Figure S5:**
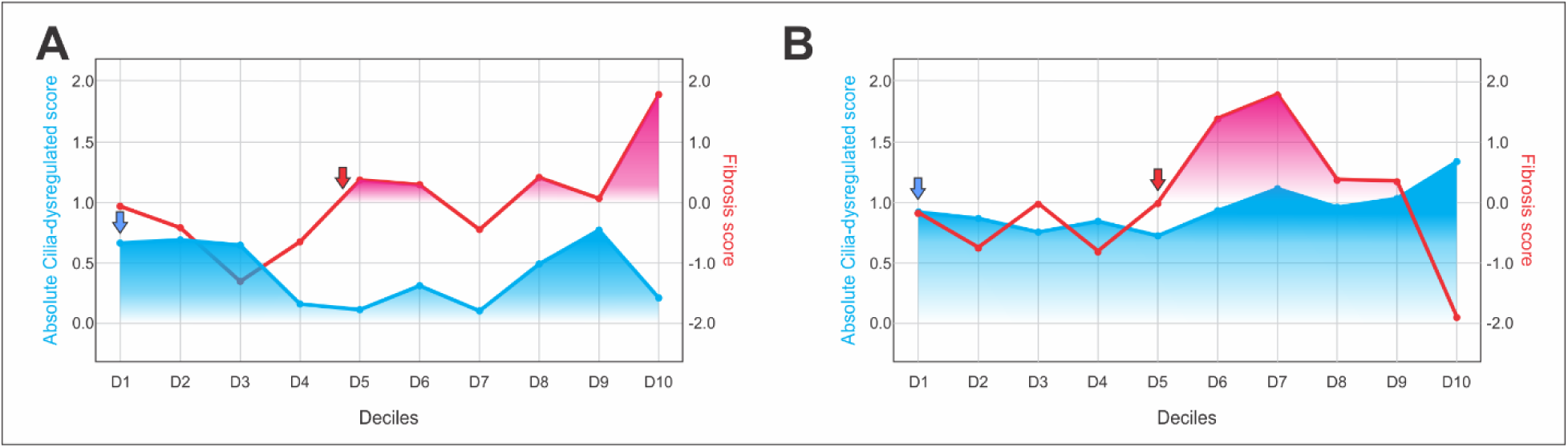
Cilia dysregulation precedes the development of fibrosis in SSc pathophysiology. Absolute cilia-dysregulated score and fibrosis score in fibroblasts from (A) Ma et al., 2024 (29) and (B) Tabib et al., 2021 (28), are shown along the pseudotime trajectory from SFRP2 healthy, SFRP2 SSc, and COL8A1 SSc fibroblasts.

**Figure S6:**
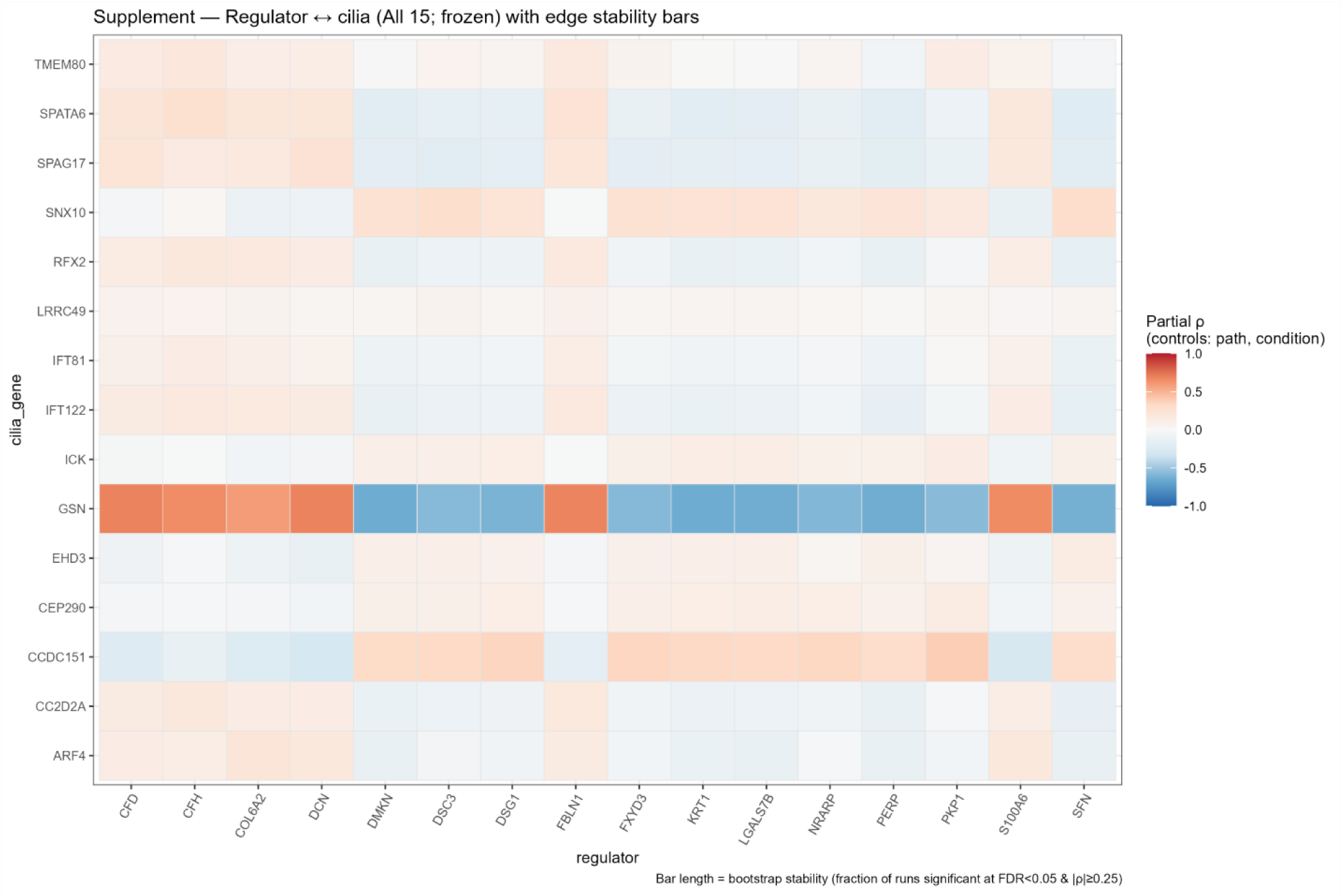
Cilia regulator genes. Partial correlation analysis identified cilia regulator modules. Cilia disassembly (high GSN) correlated positively with ECM/complement factors (DCN, FBLN1, COL6A2, CFD, CFH). In contrast, the broader cilia program correlated negatively with epidermal/keratin-junction markers (KRT1, DSG1, DSC3, PKP1, SFN, DMKN).

**Figure S7:**
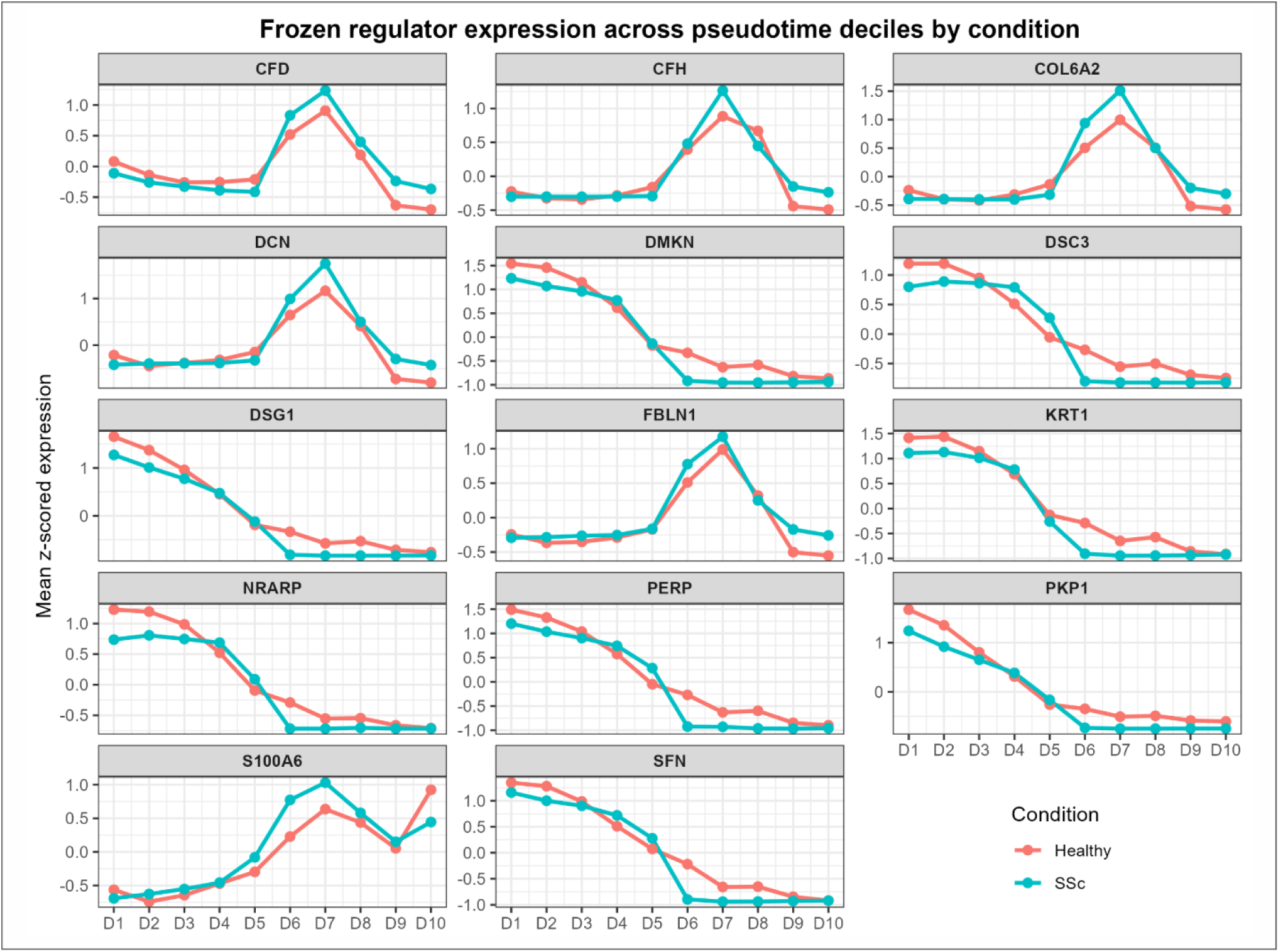
Expression of cilia regulator genes across pseudotime deciles in healthy and SSc fibroblasts. To test whether the inferred regulators were robust and not due to technical artefacts, the analysis was conducted on a decontaminated, singlet-only fibroblast subset obtained by applying DecontX (contamination ≤ 0.20) and scDblFinder to the fibroblast object with pseudotime/deciles.

**Figure S8:**
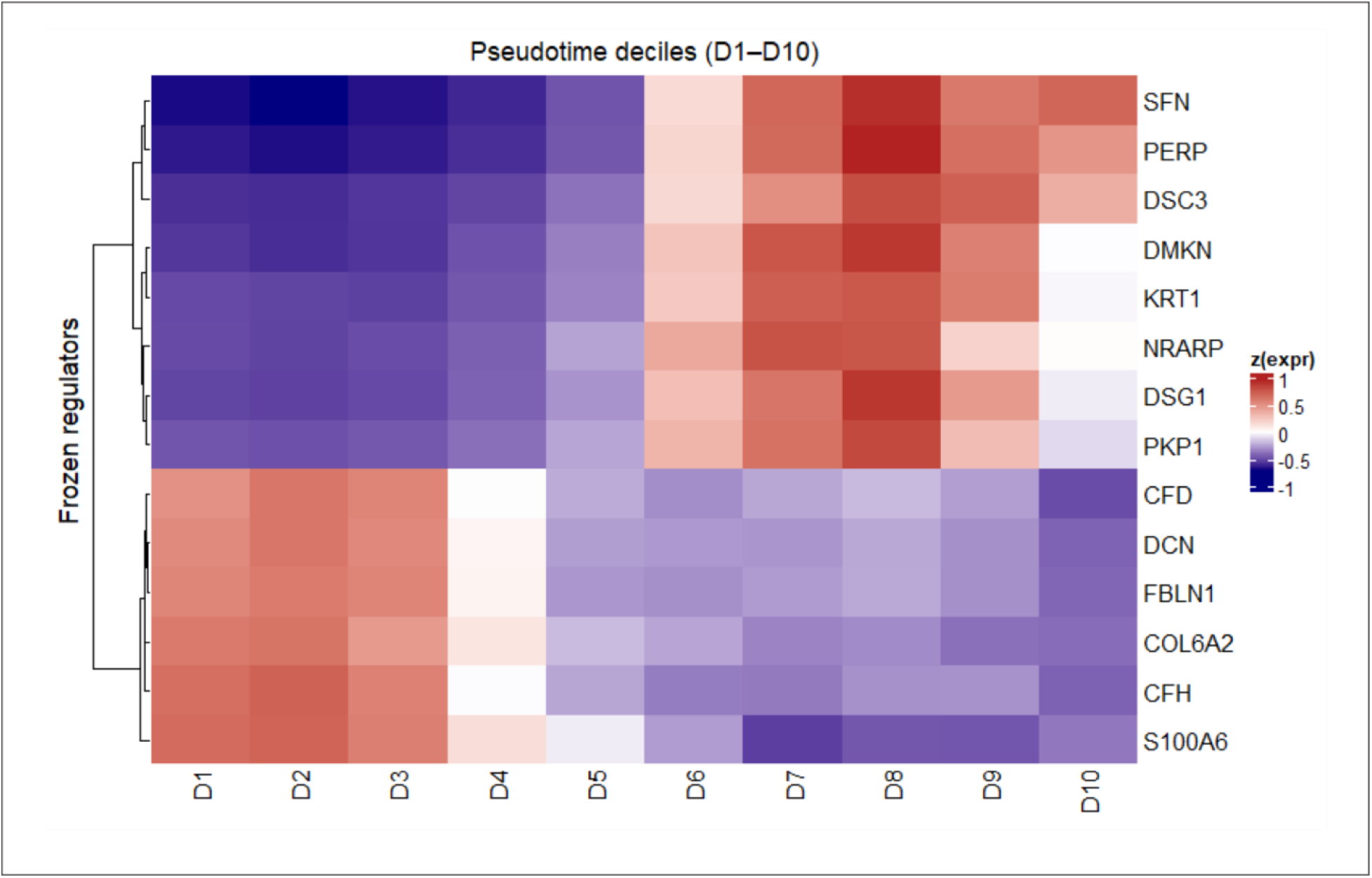
Heatmap showing the activity of 14 regulators throughout D1 to D10. Decile-wise correlation heatmaps and the QC-restricted panel confirmed that regulator trajectories are essentially unchanged after decontamination, arguing that these modules reflect genuine biology rather than doublets in a scRNAseq cluster or ambient RNA contamination.

**Figure S9:**
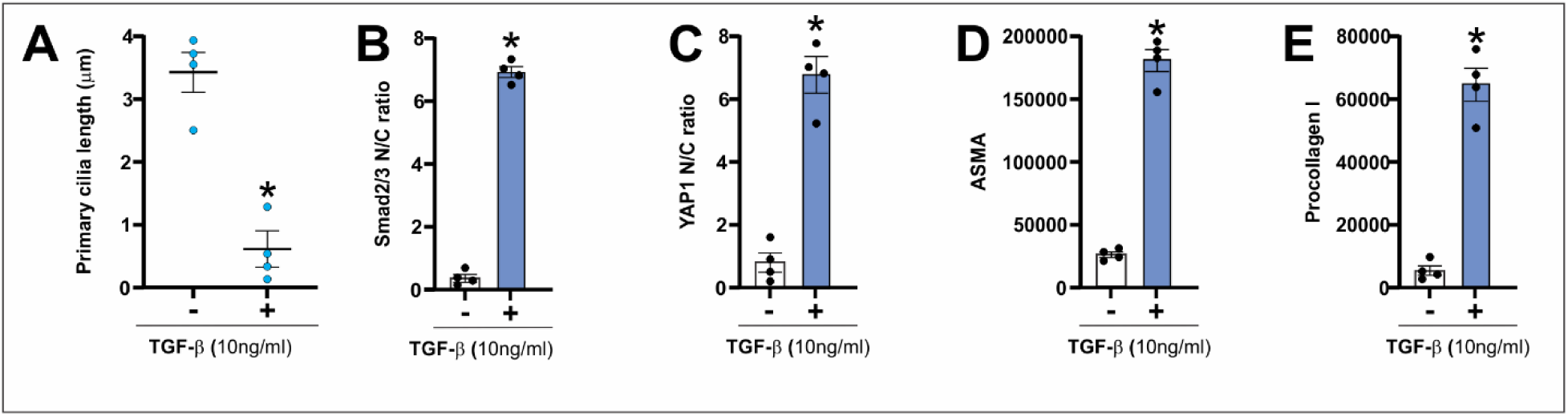
TGF-β treatment promotes shortening of PC and subsequent TGF-β and Hippo pathway activation and increase in ASMA and Procollagen. **I.** Explanted wild-type mouse dermal fibroblasts were cultured in the presence or absence of 10ng/ml TGF-β1 for 24h. Next cells were immunolabeled using anti ARL13B, ASMA, Procollagen I, SMAD2/3 and YAP1 antibodies. (A) primary cilia length, (B) SMAD2/3 N/C ratio, (C) YAP1 N/C ratio, (D) ASMA and (E) procollagen I. Statistical significance was assessed using Student’s t-test. Data are presented as mean ± SEM, n=4. * indicates statistical significance, p < 0.05.

**Table S1:**
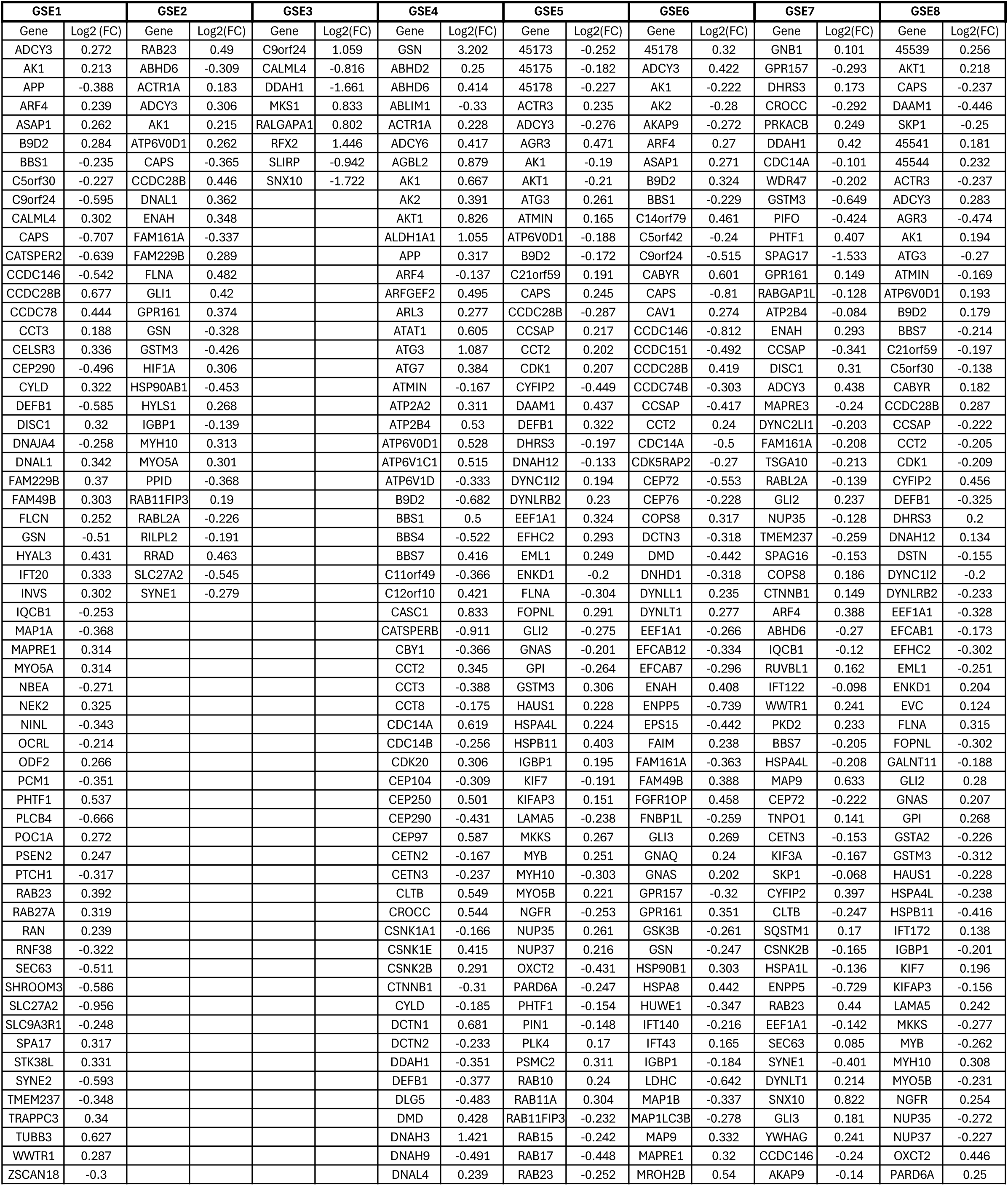
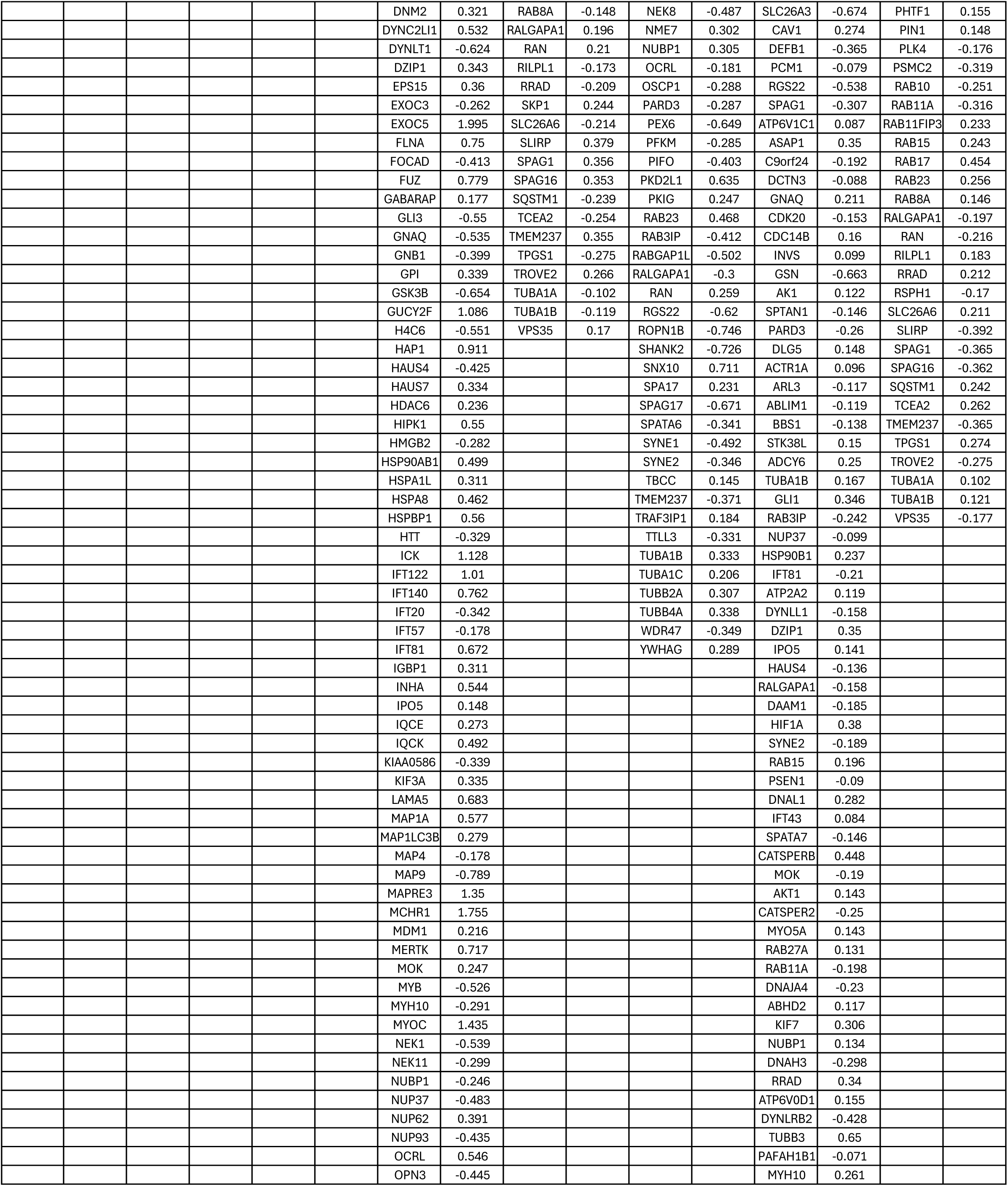
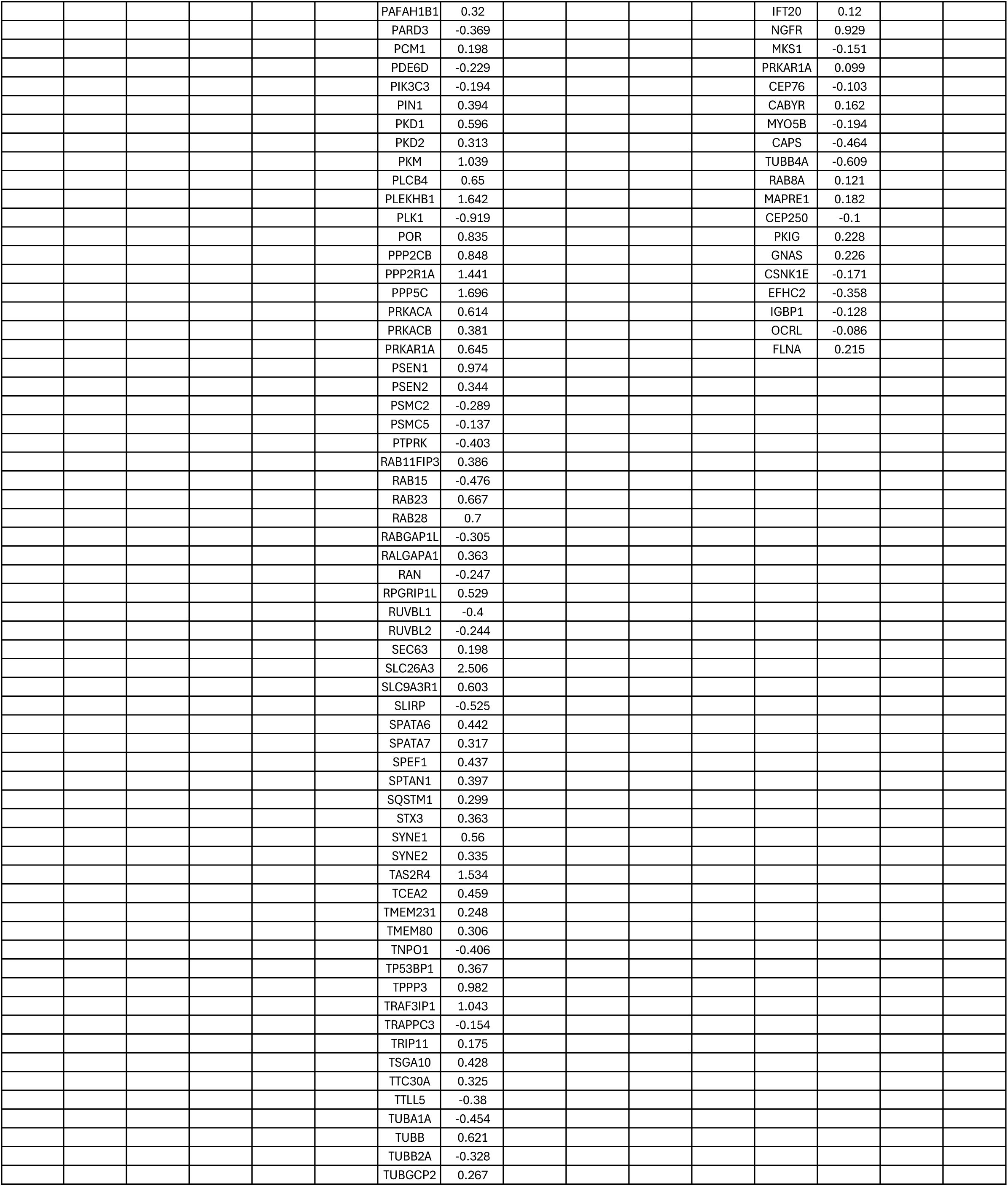
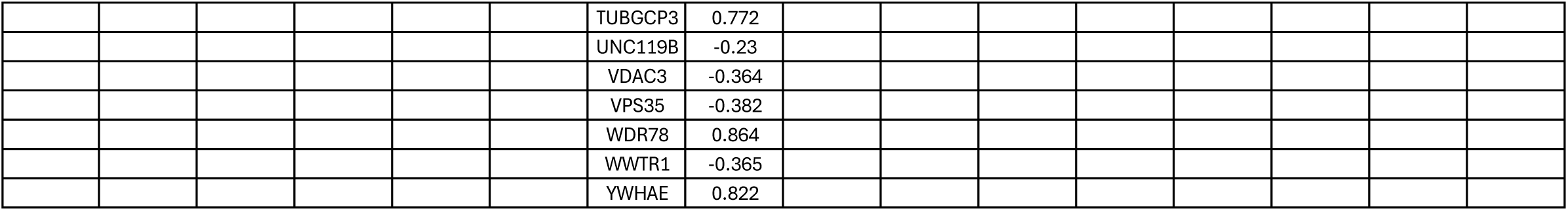
Distinct cilia-related signature recurrent in SSc across eight independent microarray datasets (GSE1–GSE8).

**Table S2:**
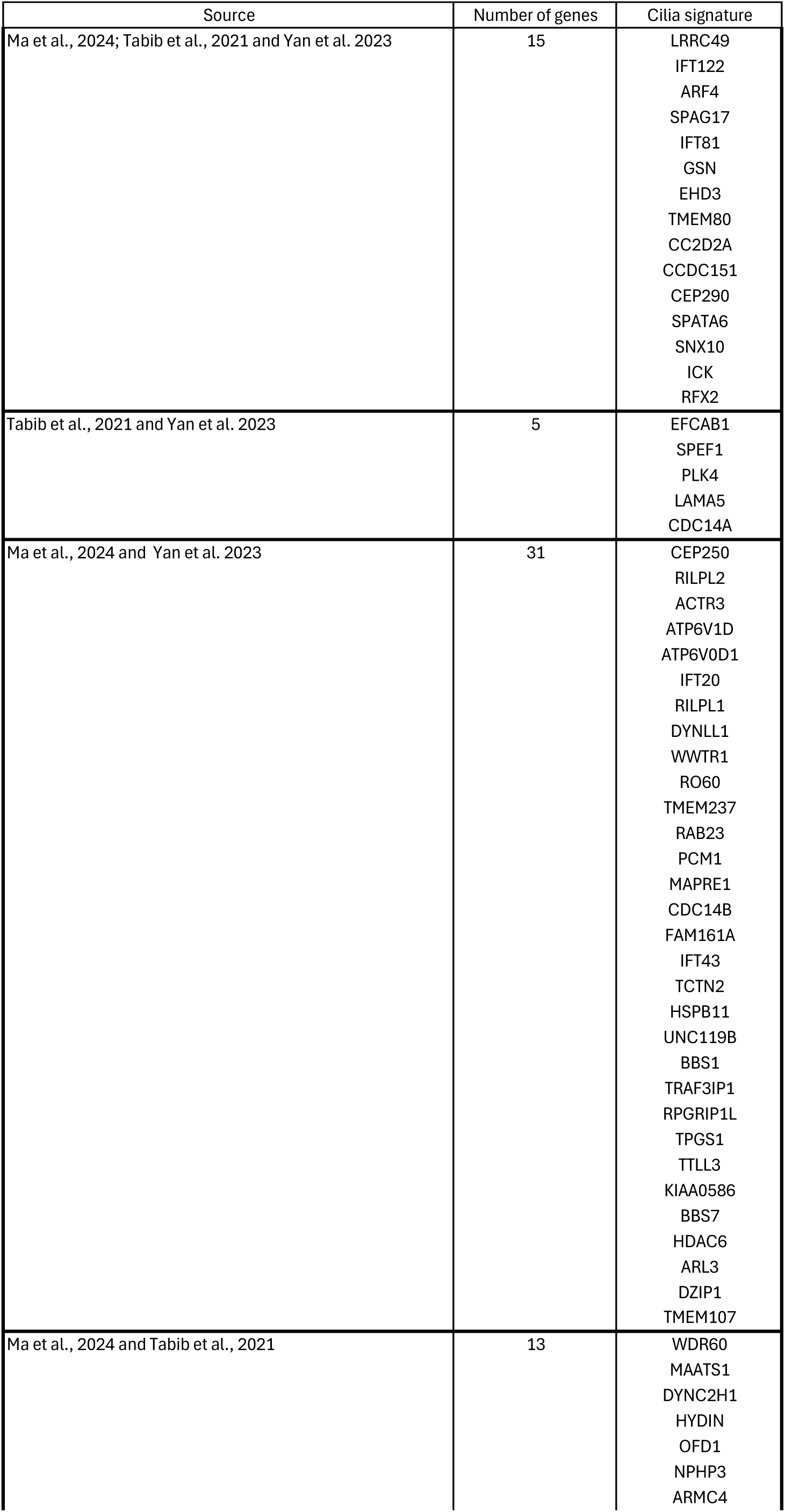
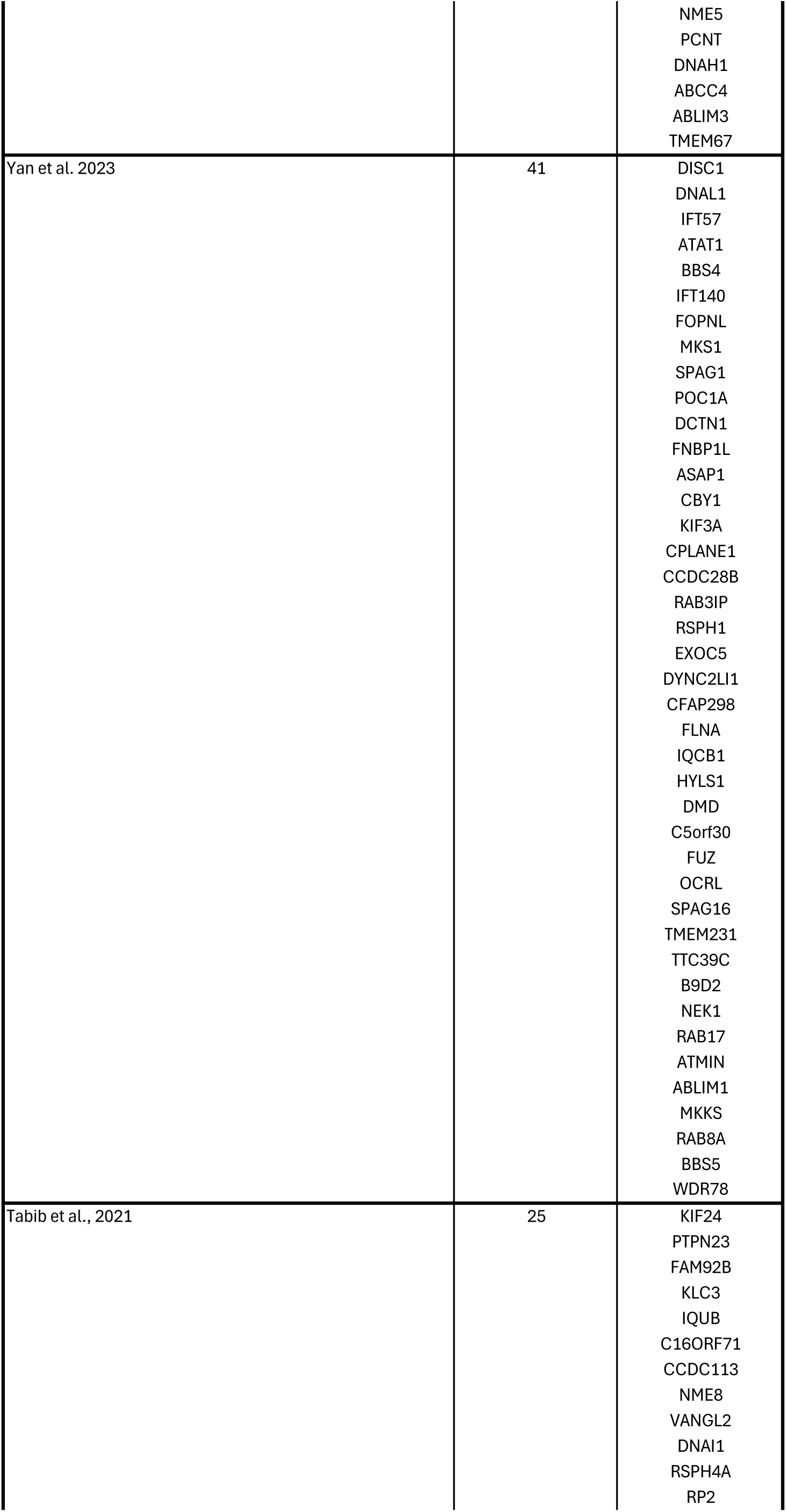
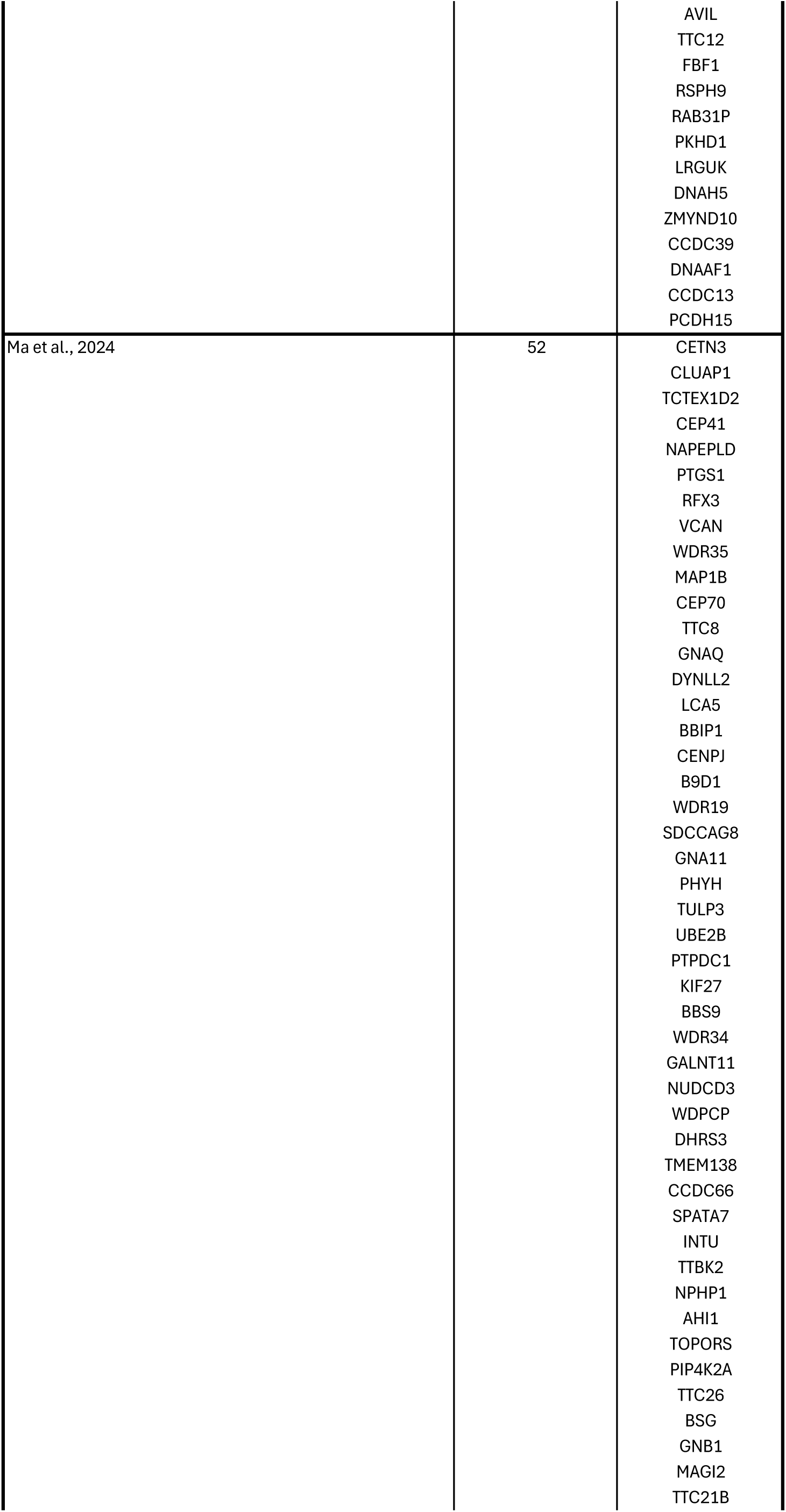
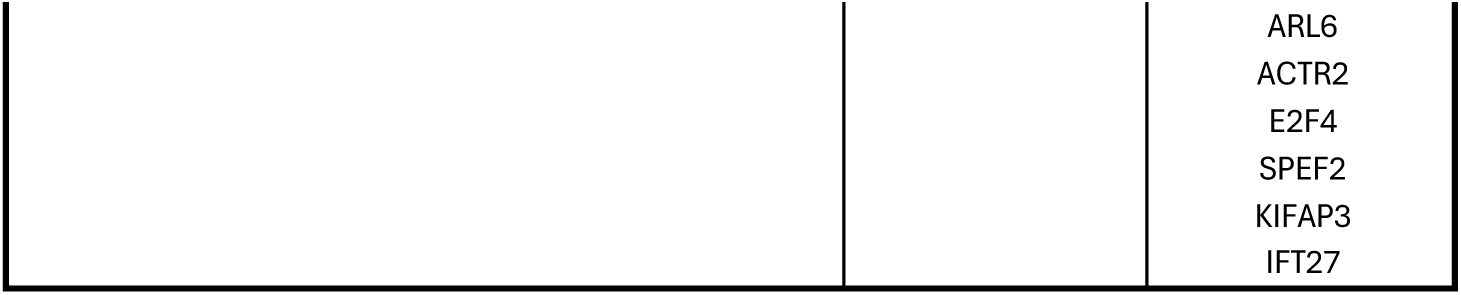
Cilia signature from Yan et al, 2023 (27), Tabib et al., 2021 (28) and Ma et al., 2024 (29).

**Table S3:** GO terms for regulators of cilia signature genes.

| Gene | GO Terms (with IDs) |
| --- | --- |
| DCN | Extracellular matrix organization (GO:0030198); Collagen fibril organization (GO:0030199); Regulation of growth factor activity (GO:0008083); Negative regulation of cell proliferation (GO:0008285); Regulation of angiogenesis (GO:0045765) |
| FBLN1 | Extracellular matrix organization (GO:0030198); Cell adhesion (GO:0007155); Basement membrane organization (GO:0071711); Elastic fiber assembly (GO:0071952); Angiogenesis (GO:0001525) |
| COL6A2 | Collagen fibril organization (GO:0030199); Extracellular matrix organization (GO:0030198); Cell adhesion (GO:0007155); Skeletal system development (GO:0001501); Response to oxidative stress (GO:0006979) |
| CFD | Complement activation, alternative pathway (GO:0006957); Proteolysis (GO:0006508); Innate immune response (GO:0045087); Positive regulation of complement activation (GO:0045957) |
| CFH | Regulation of complement activation, alternative pathway (GO:0097350); Negative regulation of immune response (GO:0050777); Innate immune response (GO:0045087); Regulation of complement-mediated cytotoxicity (GO:2000463) |
| KRT1 | Keratinization (GO:0031424); Cornification (GO:0070268); Epidermis development (GO:0008544); Keratinocyte differentiation (GO:0030216); Intermediate filament organization (GO:0045109); Response to stress (GO:0006950) |
| DSG1 | Desmosome organization (GO:0002934); Cell–cell adhesion via plasma membrane adhesion molecules (GO:0098609); Epidermis development (GO:0008544); Keratinocyte differentiation (GO:0030216); Skin barrier establishment (GO:0061436) |
| DSC3 | Desmosome assembly (GO:0002933); Cell–cell adhesion (GO:0098602); Keratinocyte differentiation (GO:0030216); Epidermis development (GO:0008544) |
| PKP1 | Desmosome assembly (GO:0002933); Regulation of cytoskeleton organization (GO:0051493); Keratinocyte adhesion (GO:0031410); Keratinocyte differentiation (GO:0030216); Epidermis morphogenesis (GO:0048730) |
| DMKN | Epidermal cell differentiation (GO:0009913); Keratinocyte differentiation (GO:0030216); Skin development (GO:0043588); Immune system process (GO:0002376) |
| SFN | Regulation of cell cycle (GO:0051726); Response to DNA damage stimulus (GO:0006974); Keratinocyte differentiation (GO:0030216); Negative regulation of cell proliferation (GO:0008285); Apoptotic signaling pathway (GO:0097190) |
| S100A6 | Regulation of cell cycle (GO:0051726); Apoptotic process (GO:0006915); Calcium ion–dependent signaling (GO:0070588); Cell proliferation (GO:0008283); Cytoskeleton organization (GO:0007010) |
| PERP | Apoptotic process (GO:0006915); Cell–cell adhesion (GO:0098602); Desmosome organization (GO:0002934); Epithelial cell differentiation (GO:0030855); Epidermis development (GO:0008544) |
| NRARP | Notch signaling pathway (GO:0007219); Negative regulation of Notch signaling (GO:0045746); Angiogenesis (GO:0001525); Cell differentiation (GO:0030154); Regulation of transcription (GO:0006355) |

